# Semaphorin 3A and 3F overexpression in TIE2 hyperactive endothelial cells contribute to the pathological lumen expansion in venous malformation

**DOI:** 10.1101/2025.07.18.665640

**Authors:** Sandra Schrenk, Chhiring Sherpa, Lindsay J. Bischoff, Yuqi Cai, Elisa Boscolo

## Abstract

**Background:** Venous malformation (VM) are developmental defects of the vasculature characterized by tremendously enlarged and dysfunctional veins. Gain-of-function somatic mutations in the endothelial-specific tyrosine kinase receptor TIE2 have been identified as leading driver of VM pathogenesis. The aim of this study was to determine whether the aberrant venous lumen expansion is caused by recruitment of wild-type endothelial cells (EC) to the lesion or by TIE2-mutant EC clonal expansion.

**Methods:** To investigate the contribution of TIE2-mutant EC and wild-type EC to the aberrant venous lumen expansion, we used a xenograft murine model of VM generated with a combination of TIE2-mutant EC and wild-type EC. To perform longitudinal studies, we employed a three-dimensional (3D) fibrin gel lumen formation assay and a migration assay, both using wild-type EC in competition or confrontation with TIE2-mutant EC. To investigate the mechanisms implicated in VM lumen expansion we used RNA-sequencing and short interference (sh)RNA in the TIE2-mutant EC.

**Results:** We demonstrate here that in the VM xenograft model, the aberrant blood vessels were lined almost exclusively by TIE2-mutant EC, and wild-type EC were rarely found. Functionally, the TIE2-mutant EC exerted a competitive advantage over wild-type EC by inhibiting wild-type EC sprouting. In line with these findings, TIE2-mutant EC promoted repulsion of wild-type EC. ShRNA-mediated silencing of Sema3A or Sema3F in TIE2-mutant EC rescued this chemorepellent phenotype and restored the ability of wild-type EC to migrate, sprout and form lumens. Furthermore, knock-down of Sema3A or 3F in TIE2-mutant EC normalized the blood vessel size in vivo.

**Conclusions:** Our results demonstrate that wild-type EC are not recruited to the aberrant veins suggesting VM pathogenesis is fueled by clonal expansion of TIE2-mutant EC. Mechanistically, we show that Sema3A and 3F are overexpressed in TIE2-mutant EC and play a crucial role in the pathological vascular lumen expansion in VM.

**Abstract (not formatted):** Venous malformations (VM) are developmental defects of the vasculature characterized by tremendously enlarged and dysfunctional veins. Gain-of function somatic mutations in the endothelial-specific tyrosine kinase receptor TIE2 have been identified as one of the leading drivers of VM pathogenesis. The aim of this study was to determine whether the aberrant venous lumen expansion is caused by the recruitment of wild-type endothelial cells (EC) to the lesion or instead by TIE2-mutant EC clonal expansion.

In a xenograft murine model of VM generated with a combination of TIE2-mutant EC and wild-type EC, we demonstrated that the aberrant blood vessels were lined almost exclusively by TIE2-mutant EC, suggesting lesions form by clonal expansion, while wild-type EC were not recruited to the aberrant veins. Functionally, in a three-dimensional cell competition assay, we showed that TIE2-mutant EC exerted a competitive advantage over wild-type EC by inhibiting wild-type EC sprouting and ability to form vascular lumens. In line with these findings, TIE2-mutant EC repelled wild-type EC by reversing their migration direction in a cell confrontation assay.

In seeking to define the mechanism driving this repulsion phenotype, we detected elevated levels of the chemorepellent Semaphorin 3A (Sema3A) and Sema3F in the TIE2-mutant EC. ShRNA-mediated silencing of Sema3A or Sema3F rescued the chemorepellent phenotype and restored the ability of wild-type EC to migrate, sprout and form lumens. Furthermore, Sema3A or 3F knock-down in TIE2-mutant EC normalized the blood vessel morphology and size in vivo. Taken together, these data strongly indicates that Sema3A and 3F are important players in VM pathogenesis.

## Introduction

Venous malformations (VMs) are developmental defects of the vasculature, characterized by abnormally enlarged veins. VMs present as blue-colored, compressible lesions and are classified as slow-flow vascular malformations, as proposed by the International Society for the Study of Vascular Anomalies (ISSVA) ^1^. The prevalence of VM is estimated to be between 1-2 in 10,000 births, occurring equally in males and females ^2^. VMs expand as the child grows, resulting in deformity, severe chronic pain, obstruction of vital organs, bleeding, thrombi, and increased risk of pulmonary embolism ^3,4^.

Standard treatments are non-targeted and invasive, often consisting of sclerotherapy alone or in combination with debulking surgery ^4–6^. Since VM is prone to relapse, patients require repeated interventions throughout their lives. There are no FDA approved medical therapies for VM. Clinical trials and pre-clinical studies ^7^ have reported that the mTOR inhibitor rapamycin can improve symptoms and suppress VM lesion expansion ^8,9^, although it does not induce significant lesion regression. Therefore, there is an urgent need to better understand the pathogenesis of VM to develop more effective and less invasive therapeutic interventions.

Gain-of-function mutations in the *TEK* gene, encoding for the endothelial tyrosine kinase receptor TIE2, have been associated with the majority (∼56%) of sporadically occurring as well as inherited cutaneous and mucosal VM ^10,11^. Somatic hyperactive mutations in *PIK3CA* (catalytic subunit of class I phosphoinositide 3-kinases), have also been associated with a smaller number of VM cases ^12,13^. TIE2 has a crucial role during vascular development as it regulates both promotion of angiogenesis and maintenance of vascular quiescence. Hyperactive mutant TIE2 and PIK3CA signaling have been shown to increase the phosphorylation of downstream pro-angiogenic pathways, including PI3K/AKT, to promote endothelial cell (EC) proliferation and survival ^7,14–17^. The most prevalent somatic mutation in VM patients is the TIE2 p.L914F (TIE2-L914F) non-synonymous variant ^10,11^.

While specific TIE2 mutations and increased AKT signaling are linked to formation and growth of VM, it is still unclear how a mutant endothelial cell or progenitor cell can drive the abnormal enlargement of the vascular lumen. An outstanding question in the field is whether mutant EC can promote cell-cell communication with neighboring wild-type EC, recruiting them to the malformed vessels, or if they can undergo clonal expansion.

Semaphorins are guidance molecules that can regulate cell-cell communication during physiological and pathological angiogenesis and endothelial cell migration. Semaphorins act by providing both repulsive and attractive signals ^18^ and they play a crucial role during the embryonic development of multiple organ systems including the nervous ^19^ and the vascular system ^20,21^. Class 3 Semaphorins are secreted guidance molecules that allow for paracrine communication. Semaphorin 3A (Sema3A) and Sema3F ^22^ exert chemorepellent actions towards EC ^21,23–25^ as well as neurons ^26–28^ by signaling through Neuropilin (NRP) and Plexin receptors ^29^. Sema3A can inhibit filopodia formation during vascular tip formation ^30^ and prevent EC integrin-mediated adhesion and vascular remodeling ^21^. While dysregulated Semaphorin class 3 expression levels are associated with pathological angiogenesis, aging-related cardiac dysfunction ^31^ and tumor progression ^32–35^, the role of Sema3A and 3F in VM pathogenesis has not yet been explored.

The aim of this study was to identify the cellular and molecular mechanisms that contribute to abnormally enlarged VM blood vessels. We thereby sought to investigate if Sema3A and 3F overexpression acts by regulating the cell-cell communication between TIE2-L914F mutant EC and wild-type EC. We employed shRNA-mediated downregulation of Sema3A or Sema3F and of their Neuropilin receptors NRP1 or NRP2 and assessed the migration of wild-type EC towards TIE2-L914F mutant EC in a cell confrontation assay, with time-lapse imaging. Next, we analyzed the role of Sema3A and 3F in the sprouting, lumen formation and expansion in a 3D in vitro cell competition assay and in a xenograft model, both obtained by intermixing TIE2-L914F mutant EC with wild-type EC.

## Methods

### Cell culture

Human umbilical vein endothelial cells (HUVEC, wild-type EC) and retrovirally transfected HUVEC expressing full-length TIE2-WT or TIE2-L914F were a gift from Dr. Lauri Eklund and Dr. Miikka Vikkula and were previously described ^7^. Endothelial Colony Forming Cells (ECFC) isolated from cord blood were a gift from Dr. Joyce Bischoff and were previously described ^36^. For co-culture assays, EC were additionally transduced with lentiviral constructs encoding for either red fluorescent protein (mCherry, Addgene, cat#36084) or green fluorescent protein (GFP) (Addgene, cat#17446-LV, ^37^).

ECs from VM patients (VM-EC) were collected from freshly resected VM lesions after informed consent as previously described ^15,38^. All the procedures were approved by the Institutional Review Board according to ethical guidelines, Approved IRB # 2016-3878 per institutional policies at Cincinnati Children’s Hospital Medical Center (CCHMC), with approval of the Committee on Clinical Investigation. Briefly, VM tissue was digested in a digestion buffer containing 1mg/mL collagenase A (Roche, cat#10103578001), and ECs were purified using CD31 immunomagnetic beads (Invitrogen, cat#11155D). Cells were cultured on fibronectin-coated (1μg/cm²) plates, in Endothelial Cell Growth Medium (EGM-2) consisting of Endothelial Cell Basal Medium (EBM-2) (Lonza, cat#CC-3156) supplemented with growth factors (EGM-2 Single Quote, Lonza, CC-4176) and 10% fetal bovine serum (FBS) (Cytiva, HyClone, cat#SH30910.03). VM patient derived EC used in this study (defined as VM#1, VM#2 and VM#3) have been characterized previously ^15^ (and correspond to patient VM9, VMA and VM11 respectively). The presence of the TIE2-L914F mutation in these cells has been confirmed by Sanger sequencing ^15^.

All cells were cultured at 37°C in a humidified 5% CO_2_ atmosphere. For conditioned media (CM) experiments, ECs were seeded at a density of 15x10^5^ cells/cm^2^. Media was collected after 48h of cell culture, centrifuged at 400 g, sterile filtered (Corning, cat# CLS431219), and used to assess wild-type EC migration or proliferation.

### Tissue Samples

The study was performed in accordance with the Declaration of Helsinki, and tissue samples from venous malformation patients’ were obtained after informed consent. All the procedures were approved by the Institutional Review Board according to ethical guidelines, Approved IRB # 2016-3878 per institutional policies at Cincinnati Children’s Hospital Medical Center (CCHMC), with approval of the Committee on Clinical Investigation. Tissue samples from venous malformation lesions were obtained from affected patients without identifiers. Each sample was further deidentified by creating a sample ID for use in this study.

### Xenograft

All animal experiments were approved by the Institutional Animal Care and Use Committee at Cincinnati Children’s Hospital (Protocol number IACUC 2023-0025). Mice were housed in the animal care facility of CCHMC under standard pathogen-free conditions with a 14 h light/10 h dark schedule and provided with food (LabDiet, cat. #5010) and water ad libitum. Xenograft studies were performed in 6–8-week-old athymic nu/nu male mice (Inotiv) as previously described ^7,38^. As the nude mice are only acting as a host for the grafted cells, using male mice that do not experience hormone cycling is the best strategy to study vasculogenesis. For the mixed xenograft model, wild-type EC (mCherry labelled – green pseudocolor) were intermixed with TIE2-L914F EC (GFP labelled – magenta pseudocolor). Cells were mixed in a ratio of 5:1 of wild-type EC:mutant EC. A total of 2.5 million cells were suspended in 200μL Matrigel® (Corning, cat#356237) and subcutaneously injected into both flanks of mice (2.09 x10^6^ WT EC and 4.1x10^5^ TIE2-L914F EC, respectively). Explants of different experimental groups were imaged from the same fixed distance, weighed, and fixed in 4% paraformaldehyde (Electron microscopy sciences, cat# 15710-S) overnight before processing for cryosectioning.

### Cryosectioning, Imaging, and Analysis

Fixed xenograft plugs were incubated in 30% sucrose solution for 24h at 4°C. Plugs were then embedded in O.C.T (optimal cutting temperature) media (Tissue-Tek^®^, cat#4583) and stored in the -80°C freezer for at least 24h prior to cryosectioning. The tissue was sectioned at 20µm and sections were stored at -20°C before being processed for imaging and analysis. Confocal z-stack images were acquired on a Nikon A1R laser scanning confocal microscope. Vessel size and vascular area (represented as percentage (%) of total tissue area) was quantified using Fiji software ^39^. For the mixed model, the number of cells lining the vessels, and the percentage of wild-type and mutant EC was calculated (as % of total number of EC lining the blood vessel).

### Immunofluorescent Staining

Immunofluorescence staining was performed on xenograft explant cryosections. Slides were blocked overnight at 4°C using 5% bovine serum albumin (BSA, Sigma, cat#A7906) in PBS (Fisher Scientific, cat#BP3994) containing 0.5% Triton X-100 (Sigma, cat#X100) and then incubated at 4°C overnight using the following antibodies: Alexa Fluor® 647 anti-mouse TER-119 (BioLegend, cat#116218, 5µg/mL) or Ulex Europaeus Agglutinin I (UEA I, 1µg/mL), DyLight® 594 (Vector Laboratories, DL-1067-1, 1µg/mL). Samples were counterstained with 4’,6-Diamidino-2-Phenylindole, Dihydrochloride (DAPI, cat#D1306, Invitrogen, 5 µg/mL) for 5 min at room temperature (RT). Sections were mounted with Fluoromount-G® mounting medium (Southern Biotech, cat#0100-01).

### RNA Scope

For the RNAscope assay, patient tissue sections were deparaffinized and endogenous peroxidase was quenched by RNAscope Hydrogen Peroxide Solution (Advanced Cell Diagnostics (ACD), cat#322381) for 10min. Samples were treated with RNAscope Target Retrieval reagent (ACD, cat#322000) for 15min at 110°C using a Decloaking Chamber (Biocare Medical). Slides were transferred into fresh 100% EtOH for 3min and air dried for 5min. Samples were incubated with 2 drops of RNAscope Protease Plus (ACD, cat#322381) for 10min at 40°C. Sections were incubated with the following probes for 2h at 40°C: C1 RNAscope® Probe - Hs-SEMA3F (Advanced Cell Diagnostics, cat#432201) and was detected with TSA Vivid™ Dyes 650 fluorophore (ACD, cat#323273, 1:750). C2 probe labeled Hs-SEMA3A (Advanced Cell Diagnostics, cat#416561-C2) and was detected with the TSA Vivid™ Dyes 570 fluorophore (ACD, cat#323272,1:750). After probe hybridization, the samples were processed for Ulex Europaeus Agglutinin I (UEA1) fluorescence labelling to identify endothelial cells. Endogenous biotin and streptavidin binding sites were blocked using the Streptavidin/Biotin Blocking Kit according to manufacturer’s protocol (Vector laboratories, cat#SP-2002). Slides were then blocked for 30min at RT using the Carbo-Free™ Blocking Solution (Vector laboratories, SP-5040-125) and incubated with biotinylated UEA1 (Vector laboratories, cat#B-1065-2, 4µg/mL (1:500) in PBS) for 1h at RT. Slides were incubated with a fluorescent Streptavidin, DyLight® 488 (SA-5488-1). Counterstaining was performed by using DAPI (ACD, cat#323108) and slides were mounted using ProLong™ Gold Antifade Mountant (Invitrogen, cat#P36930).

### 3D cell competition assay

The 3D cell competition assay was adapted from the 3D fibrin bead lumen formation assay previously described ^40,41^. Briefly, Cytodex® 3 microcarrier beads (Sigma, cat#C3275) were separately coated with either wild-type EC or TIE2-L914F EC (fluorescently labelled) at a concentration of 400 cells/bead for 4h at 37°C. The following day, coated beads were mixed at a ratio of 1:1 (wild-type EC coated beads:mutant EC coated beads) and resuspended in 2mg/mL of fibrinogen (Sigma, cat#F8630) solution containing 0.15U/mL of aprotinin (Sigma, cat#A1153) at a concentration of 500 beads/mL. Then 0.625U/mL of thrombin (Sigma, cat#T4648) and 0.5mL beads/fibrinogen suspension were carefully added to a well of a 24-well plate and incubated at 37°C for 20min to allow fibrin clotting. The gels were overlaid with human dermal fibroblasts (ATCC, cat#PCS-201-012) at 2×10^4^ cells/well and cultured in EGM2 supplemented with 10% FBS. Medium was replaced every other day. Images were acquired with EVOS (Invitrogen) or Nikon A1R laser scanning confocal microscope at the indicated time points, and the number of sprouts per bead were counted manually. Z-stack images were reconstructed with Imaris software (Oxford instruments).

### Proliferation analysis

For proliferation analysis, EC treated with either control Vector, shSema3F, or shSema3A were plated onto 96 well plates at 6000 cells/well and cell growth was observed with 12 replicates per condition. For proliferation analysis with conditioned medium, wild-type EC were plated onto 96-well plates at 6000 cells per well. Wild-type EC were then incubated with conditioned medium (collected from wild-type EC or TIE2-L914F EC) with 8 replicates per condition. Cell growth was monitored for 3 days using an IncuCyte ZOOM Live Cell Imaging System (Essen BioScience).

### Transwell migration assay

For Transwell migration assay, 5 × 10^4^ cells were seeded into the cell culture insert (upper chamber of the 8-μm pore size membrane insert) (Corning Costar, cat# CLS3464) and allowed to settle for 1h. Conditioned Medium (wild-type, TIE2-WT or TIE2-L914F EC conditioned for 48h as described above) was added to the lower chamber. After 6h of incubation, the non-migrated cells remaining in the upper membrane were removed with cotton applicators. The cells that migrated through the membrane were fixed and stained with 1% crystal violet solution (Millipore #V5265), imaged and counted under an EVOS cell imaging system (Invitrogen) and shown as migrated wild-type EC per field.

### Scratch wound healing assay

Cell migration was assessed using a scratch wound healing assay ^42^. In brief, cells were cultured in 24-well plates at density of 2×10^5^ cells/cm^2^ and grown to confluency before a scratch was created using a 20µl pipet tip. Cells were washed with PBS to remove any debris, and the medium was replaced with conditioned medium (wild-type, TIE2-WT or TIE2-L914F EC conditioned for 48h as described above) containing 2mM hydroxyurea to prevent cell proliferation. Phase-contrast images were captured at 0h and 20h after wounding using an EVOS-imaging system (Invitrogen). At the endpoint, cells were fixed in 4% PFA (Electron microscopy sciences, cat# 15710-S) for 15min at RT and visualized using Alexa Fluor 488 Phalloidin (Thermo Fisher, cat# A12379) on an EVOS. The gap (cell-free) area was measured using Fiji software, and the value was expressed as percentage of the initial gap area at 0h (% gap closure area).

### Cell confrontation assay

For cell confrontation assays, a two-well culture insert (Ibidi, cat# 80209) was placed onto a glass bottom 24-Well plate (Cellvis, cat#P24-1.5H-N). These silicone inserts create a gap via cell exclusion by preventing cell growth in between two wells and create a reproducible gap width of about 500µm. 2000 cells were seeded into each compartment and cells were allowed to grow to confluency and the cell migration into the gap area was initiated by removing the silicone insert. Images of the same gap area were captured every 30min for a total of 15h using a Nikon Ti2 Widefield microscope at 37°C with 5% CO_2_. Cell nuclei were visualized with NucBlue™ Live ReadyProbes™ Reagent (Thermo Fisher, cat#R37605). Individual cells (≥ 20 cells/condition) were tracked over time using manual cell tracking plugin (https://imagej.net/ij/plugins/track/track.html) for Fiji software. Directionality, forward migration index, velocity, euclidean distance, and accumulated distance, as well as rose plots, were calculated using the Chemotaxis and Migration Tool (Ibidi, https://ibidi.com/chemotaxis-analysis/172-manual-tracking.html).

### RNA Sequencing

Total RNA was isolated from WT EC, TIE2-WT and TIE2-L914F EC (n=4 biological replicates/group) using RNeasy extraction kit (Qiagen, cat#74106) according to manufacturer’s recommendations. The quality of RNA was analyzed using a Bioanalyzer (Agilent), which showed high quality (RIN score >9.5). The directional polyA RNA-seq was performed by the Genomics, Epigenomics and Sequencing Core at the University of Cincinnati using established protocols as previously described ^43,44^. To enrich polyA RNA for library preparation, NEBNext Poly(A) mRNA Magnetic Isolation Module (New England BioLabs) was used with 500 ng total RNA as input. Next, NEBNext Ultra II Directional RNA Library Prep kit (New England BioLabs) was used for unique dual index library preparation under PCR cycle number of 9. After library QC and Qubit quantification (ThermoFisher), the normalized libraries were sequenced using NextSeq 2000 Sequencer (Illumina). Under the setting of PE 2x61 bp, ∼30M read pairs were generated for each sample. Fastq files for downstream data analysis were automatically generated via the Illumina BaseSpace Sequence Hub after sequencing. Differential gene expressions between groups (TIE2-WT EC vs TIE2-L914F EC) of n=4 samples/group were assessed with the R package DESeq2 ^45^. The log2 expression fold-change threshold of |0.58| and FDR (false discovery rate) adjusted P-value < 0.05 were used to identify DEGs. Gene ontology (GO) analyses were performed using the enrichment analysis tool Enrichr ^46–48^.

### Real time PCR

Reverse transcription was performed using an iScript cDNA synthesis kit (Bio-Rad, cat#1708841). qPCR was performed using SsoAdvanced Universal SYBR(R) Green Supermix (Bio-Rad, cat#1725272). Amplification was done in a Bio-Rad Touch Real-time PCR detection system (Bio-Rad, cat#CFX96). A relative standard curve of each gene amplification was generated and an amplification efficiency of >90% was considered acceptable. Hypoxanthine phosphoribosyl transferase 1 (*HPRT1*) and TATA-binding protein 1 (*TBP1*) were used as housekeeping genes. Quantification was performed using the Pfaffl method ^49^. Primer sequences are shown in Supplementary Table 1.

### Western Blot and ELISA assay

Cells were lysed using ice-cold radioimmunoprecipitation assay (RIPA) lysis buffer (Boston Bioproduct) containing HALT^TM^ protease/phosphatase inhibitor cocktail (Thermo Scientific, cat# 78442). The protein concentration was determined using the BCA Protein Assay Kit (Thermo Fisher, cat#PI23225). 20-40 µg of total protein were subjected to SDS-PAGE (4-20%, Midi Criterion precast gels, Bio-Rad, cat#5678094) and transferred to a PVDF membrane (Immobilon®-P PVDF Membrane, Millipore, cat#IPVH00010). Membranes were blocked for in 5% nonfat dried milk for 1h at RT and probed with the following antibodies: anti-Sema3A (Sigma, cat#AB9604,1 µg/mL), anti-Sema3F (Sigma, cat#AB5471P, 1 µg/mL), anti-Neuropilin-1 (Cell signaling, cat#54568, 1:1000 dilution), anti-Neuropilin-2 (Invitrogen, cat#PA5-65910, 0.2µg/mL) and anti-GAPDH (Sigma, cat#MAB374, clone 6C5, 0.2 µg/mL) overnight at 4°C. Membranes were incubated with anti-mouse or anti-rabbit Peroxidase conjugated secondary antibodies for 1h at RT. All washes were performed using TBS with 0.5% Tween 20 (Bio-Rad, cat#1706531). Antigen–antibody complexes were visualized using Immobilon Classico Western HRP substrate (Millipore, cat# WBLUC0500) and a ChemiDoc Imaging system (Bio-Rad, cat#12003153). Sema3A and Sema3F protein levels were quantified using a total of 20ug of protein lysate. The following ELISA assays were utilized according to manufacturer’s instructions: Sema 3A (Invitrogen, cat# EEL174), Sema3F (Creative Diagnostics, cat# DEIA-FN1341).

### shRNA-mediated gene knock-down

For knock-down experiments, we used MISSION® shRNA lentiviral Transduction particles (pLKO.1) which include a puromycin selection marker (Sigma): shSema3F (TRCN0000061630), shSema3A (TRCN0000058141), shNRP1 (TRCN000006327), shNRP2 (TRCN0000063312), and Control Vector (SHC001, SHC016V) (all purchased from Sigma). For transduction we followed the manufacturer’s instructions. Briefly, subconfluent EC monolayers were transduced with the lentivirus at an MOI of 5. The next day, puromycin was added to the media at a concentration of 1μg/mL to select for cells expressing the construct. After two passages, each shRNA sequence was then tested for the degree of knock-down by real-time PCR and Western Blotting or ELISA Assay.

### Statistical Analysis

Data analyzed statistically are presented as mean±standard deviation (SD) of two or more biological replicates (*n* values reported in figure legends). Statistical significance between two groups was assessed by parametric Welch’s t-test. When more than two groups were compared, one or two-way ANOVA followed by Dunnett’s or Sidak’s post hoc test was used. All calculations were performed using GraphPad Prism. Differences were considered significant for *P* value ≤ 0.05.

## Results

### TIE2-L914F EC do not recruit wild-type EC in the ectatic lumens in vivo and inhibit 3D morphogenesis of wild-type EC

Endothelial cells engineered to express the hyperactive TIE2 p.L914F mutant receptor are known to exhibit ligand-independent phosphorylation of TIE2 and increased downstream activation of the AKT pathway^50,51^, a result we confirmed in our cells (Suppl. Fig. 1a). When these EC harboring the hyperactive TIE2 p.L914F mutation are injected into the flanks of immunocompromised nude mice, they generate lesions with ectatic blood vessels, mimicking the pathological lesions of VM patients ^7,38^. To understand the processes involved in the formation of VM lesions and investigate whether wild-type EC are recruited into lesional vessels, we used a modified version of this xenograft model. We intermixed fluorescently labeled wild-type EC (shown in green) and TIE2-L914F EC (shown in magenta) and injected them subcutaneously into mice, within one xenograft plug (Fig. 1a). To account for the proliferative advantage of TIE2-L914F EC ^41^, cells were injected in a ratio of 5:1 (wild-type EC: TIE2-L914F EC). Xenograft plugs were analyzed macroscopically on day 14 after cell injection. All the plugs were visually vascularized and perfused (Fig. 1b, Suppl. Fig. 1b-c). Histologically, the enlarged vascular lumens were lined almost exclusively by TIE2-mutant EC (98.3%), while only a small percentage of wild-type EC were found in the lesional vessels (1.7%) (Fig. 1c-d). These data show that only rare wild-type EC are incorporated into aberrant veins.

**Figure 1.**
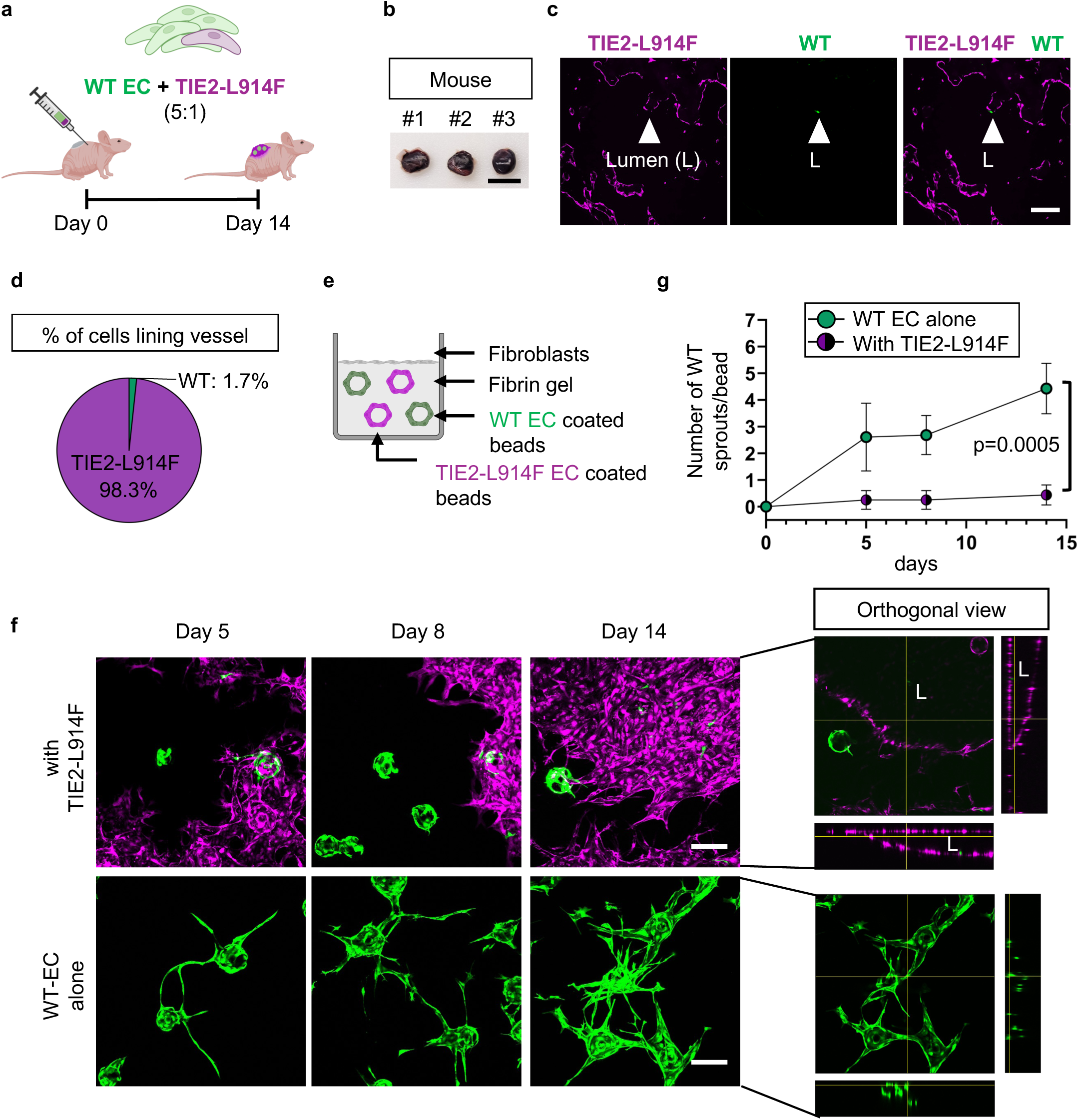
Wild-type EC are rarely found in aberrantly enlarged vessels in a xenograft model of VM. (a) Schematic of the mixed EC xenograft model of VM: wild-type (WT) EC (HUVEC, in green) were intermixed with TIE2-L914F EC (HUVEC TIE2-L914F, in magenta) at a 5:1 ratio and injected in the flanks of immunocompromised nude mice. (b) Photograph of representative xenograft explants at day 14 after cell injection. Scale bar: 1cm. (c) Tissue sections of the xenograft explant showing TIE2-L914F mutant EC (shown in magenta), and wild-type EC (shown in green) lining the vascular lesion (arrowhead). Scale bar: 200µm (d) Quantification of the cell types lining the ectatic blood vessels shown as % of cells lining vessel. n=7 mice and n=122 vessels. Mean±SD, Welch’s t-test. P-values are indicated. (e) Schematic of 3D cell competition assay: dextran beads are separately coated with TIE2-mutant EC (magenta) or wild-type EC (green) and then combined in the 3D fibrin gel (1:1 ratio). (f) Confocal z-stack images (max. intensity projection) of wild-type EC alone (bottom) or combined with TIE2-mutant EC (top) at day 5, 8 and 14. L: Lumen. Scale bar: 200µm. (g) Quantification of wild-type EC sprout number per bead. Mean±SD, 2-way ANOVA. p-values are indicated. Schematic in (a, e) were created with Biorender.com.

When wild-type EC grow within 3-dimensional (3D) fibrin gels, they generate thin, extended vascular sprouts with hollow lumens. Conversely, TIE2-mutant EC form ectatic, cyst-like channels that expand over time, invading the gel ^40,41^. To investigate if TIE2-L914F EC affect wild-type EC morphogenesis, we established a ‘3D cell competition assay’. In this assay, we coated dextran beads with fluorescently labeled wild-type EC (shown in green) and separately with TIE2-L914F EC (shown in magenta). Next, these EC-coated beads were embedded in the fibrin gel together (Fig. 1e) and topped with fibroblasts. The sprouting/tube formation behavior of wild-type EC was analyzed over a time course of 14 days. Noticeably, when in presence of the TIE2-mutant EC, the sprouting ability of wild-type EC was greatly reduced, and lumens did not form (Fig. 1f). Quantification revealed that in the presence of TIE2-L914F EC, the number of wild-type EC sprouts per bead was significantly decreased compared to native sprouting behavior and remained stagnant over the time course of 14 days (Fig.1g).

Taken together, these in vivo and 3D in vitro data indicate that TIE2-mutant EC have a competitive advantage over wild-type EC. We define here ‘competitive advantage’ as a process in which TIE2-mutant EC form enlarged lumens while inhibiting wild-type EC morphogenesis.

### TIE2-L914F EC inhibit migration of wild-type EC

Migration and proliferation are the first events during morphogenesis that lead to sprouting and new vessel formation in the 3D fibrin gel assay ^52,53^. Based on these results, we hypothesized that TIE2-L914F EC can exert a competitive advantage over wild-type EC migration and/or proliferation by means of secreted factors.

Using a Transwell migration assay we assessed the ability of secreted factors from TIE2 mutant EC to influence wild-type EC migration towards a concentration gradient of secreted factors (chemotaxis). Wild-type EC were seeded onto the top chamber of a Transwell and then incubated in conditioned medium (CM) from wild-type or TIE2-L914F EC for 6h (Fig.2a). Quantification of wild-type EC that migrated across the Transwell towards TIE2-L914F CM showed a significant (p<0.001) reduction compared to the migration towards CM from wild-type EC (Fig.2b-c). Furthermore, a scratch-wound assay was performed on confluent wild-type EC monolayers challenged with CM (Suppl. Fig. 2a). To make sure that migration differences were not confounded by changes in proliferation, the monolayers were pretreated with 4-hydroxyurea to inhibit cell division during the migration period. CM from TIE2-L914F EC induced a significant (p=0.0045) reduction of wild-type EC wound closure over a 20h period compared to that obtained with the use of CM from wild-type EC or TIE2-WT EC (Suppl. Fig. 2b-c). Next, the effect of TIE2-L914F EC CM on cell proliferation of wild-type EC was analyzed using a live cell imaging assay which revealed no significant inhibition of wild-type EC proliferation (Suppl. Fig. 2d).

Based on these results, we performed more in-depth analysis of cell migration behavior. Thereby, we performed a ‘cell confrontation assay’ by seeding TIE2-mutant and wild-type EC opposite (confronting) of each other (Fig. 2d). The migration of wild-type EC towards TIE2-mutant EC was visualized through time-lapse imaging over a time course of 15 h. Individual cells were identified by their nuclei, and their movement was tracked over time (Fig. 2e). The cell tracking analysis was divided into a) initial migration phase (0-10 h) and b) confrontation phase (10-15 h) of the assay. During the initial migration phase wild-type EC in the control setup (wild-type EC alone) migrated in a direct and straightforward pattern into the gap, while their migration pattern towards the TIE2-mutant EC was more disorganized (Fig.2f, left panel).

**Figure 2.**
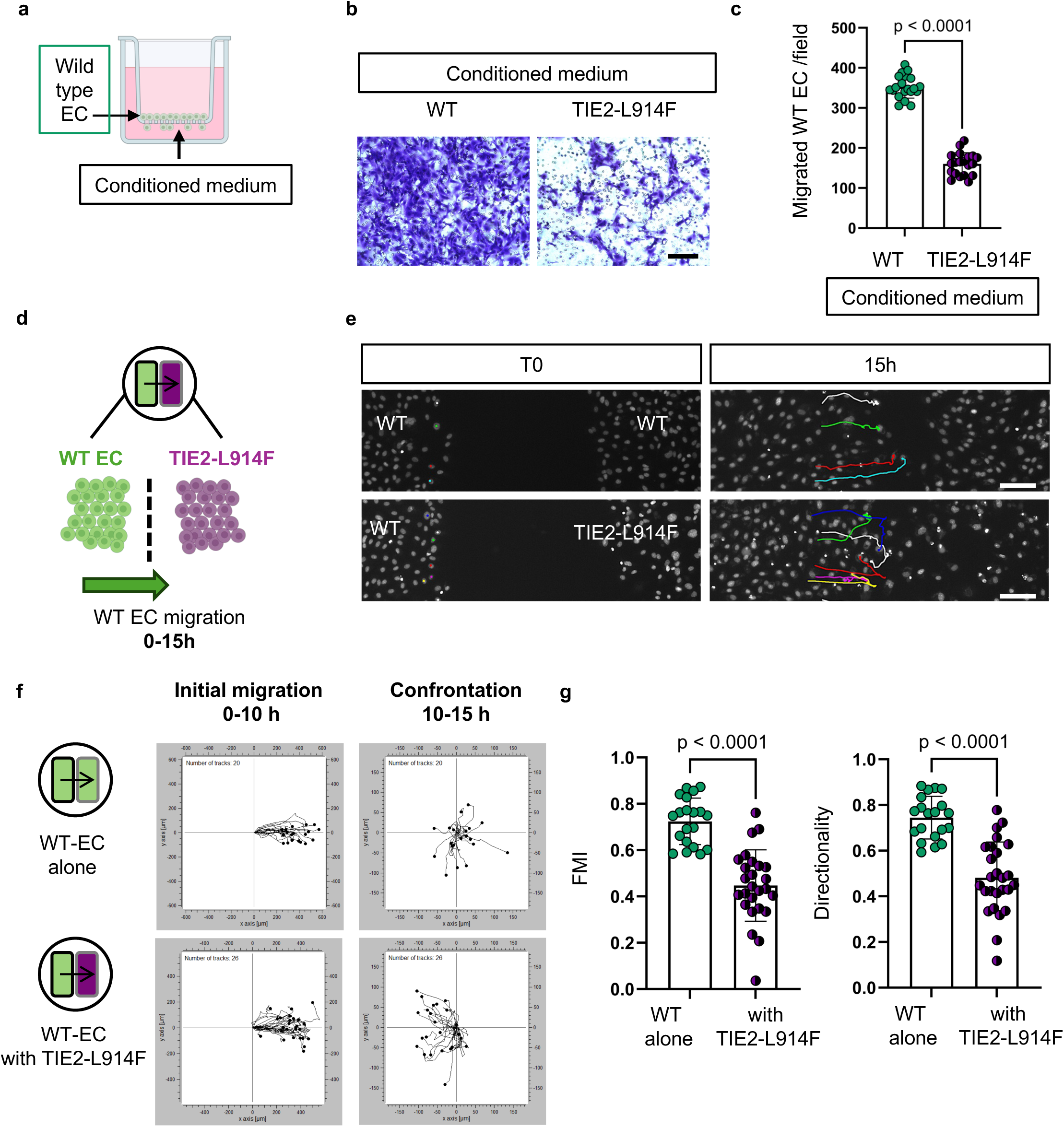
Mutant EC repulse wild-type EC. (a) Schematic of transwell migration assay. Schematic was created with Biorender.com. (b) Representative images of migrated wild-type cells towards wild-type or TIE2-L914F EC conditioned medium after 6h of migration. (c) Quantification of number of migrated wild-type cells per field. n=20 fields from n=2 membranes were imaged and quantified. Mean±SD, Welch’s t-test. P-values are indicated. (d) Schematic of cell confrontation assay. (e) The migration of wild-type EC toward TIE2-mutant EC was visualized through time-lapse imaging and then analyzed via cell tracking. Each individual cell is represented by a different color. Scale bar: 100µm (f) Trajectory plots of single cell movements in the initial migration (0-10h) and confrontation phase (10-15h). (g) Quantification of forward migration index (FMI) and directionality during the whole duration of the assay (0-15h). n≥20 cells per condition were tracked. Mean±SD, Welch’s t-test. P-values are indicated.

During the confrontation phase, wild-type EC in the control setup showed random patterns of motion, similar to what is observed in confluent or semi-confluent cell monolayers (Fig. 2f, upper right panel). Surprisingly, when confronted with TIE2-mutant EC, the wild-type EC inverted their migration direction, indicating they were repulsed away by the mutant EC (Fig. 2f, lower right panel).

This repulsive mechanism exerted by TIE2-mutant EC towards wild-type EC is demonstrated by a marked reduction in the forward migration index (FMI) (a measure of efficiency of forward migration) (Fig. 2g) and in the directionality (a measure of straightness of a trajectory). Additionally, the wild-type EC in confrontation with TIE2-mutant EC showed an increase in velocity and total accumulated distance, while the Euclidean distance (the direct distance from start point to end point) was shorter (Suppl. Fig. 3), confirming the rather disorganized and inefficient migration behavior when compared to the control setup.

### Chemorepellent Semaphorin 3A and 3F are overexpressed in TIE2-mutant EC

Next, we sought to identify the paracrine signals involved in the repulsive activity of TIE2-L914F EC on wild-type EC migration. To identify potential candidates, we performed RNA sequencing analysis comparing the gene expression of TIE2-L914F, TIE2-WT and WT EC. Principal component analysis (PCA) showed a clear separation of gene expression between TIE2-L914F EC, TIE2-WT and wild-type EC (Suppl. Fig. 4a). RNA sequencing analysis of TIE2-mutant EC, when compared to wild-type EC and EC expressing wild-type-TIE2 (TIE2-WT EC), revealed a total of 1978 differentially expressed genes (DEGs). A total of 860 genes showed increased expression while 1118 genes showed decreased expression (Fig. 3a). Gene ontology analysis showed an upregulation of the following molecular functions: Chemorepellent activity, Neuropilin binding, and Semaphorin receptor binding (Fig. 3b) – which were among the top 25. This suggested that Semaphorins, guidance molecules that are implicated in regulating cell-cell communication in the vascular system, may be involved in the chemorepellent properties of TIE2-L914F EC ^25,26^. Real time PCR (Suppl. Fig.4b), immunoblotting analysis (Fig. 3c), and ELISA assay (Fig. 3d-e) confirmed that TIE2-L914F EC expressed significantly (p<0.05) elevated levels of the chemorepellent molecules Semaphorin 3A (Sema3A) and Sema3F.

**Figure 3.**
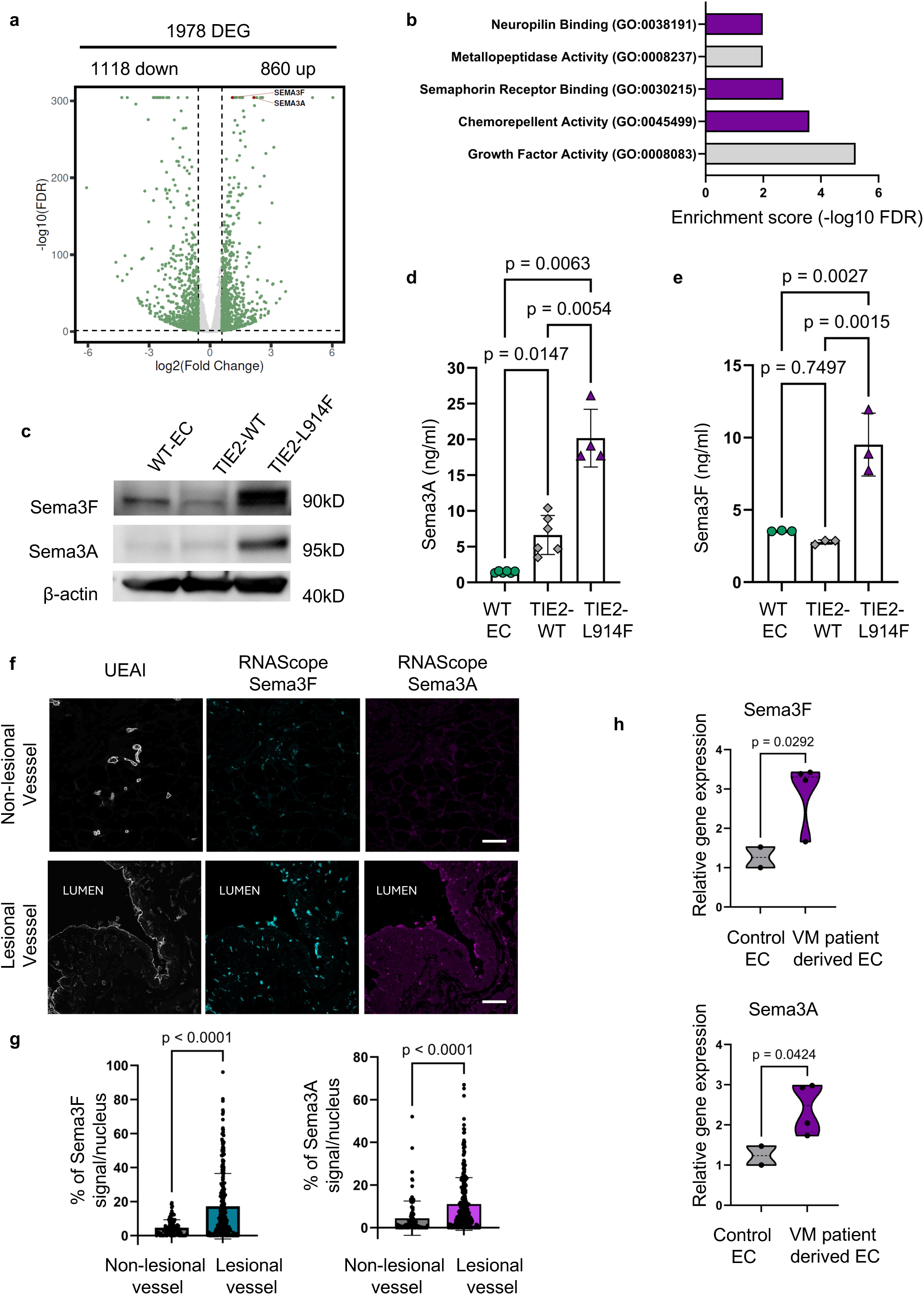
Chemorepellent Sema3A/3F are overexpressed in TIE2-mutant-EC. (a) Volcano plot of DEGs in TIE2-L914F compared to TIE2-WT EC. A total of 860 genes showed increased expression while 1118 genes showed decreased expression. Upregulated gene expression of Sema3A and Sema3F are highlighted. (b) Gene ontology analysis of RNA sequencing of TIE2-mutant EC compared to TIE2-WT EC. In TIE2-mutant EC the following molecular functions were amongst the top 25 upregulated: Chemorepellent activity, Semaphorin receptor binding and Neuropilin binding. (c) Immmunoblot of Sema3F and Sema3A in WT-EC, TIE2-WT and TIE2-L914F EC. (d) Quantitative ELISA assay of Sema3A and (e) Sema3F protein levels in WT-EC, TIE2-WT and TIE2-L914F cells. Mean±SD, one-way ANOVA, p-values are indicated. (f) RNA Scope assay showing Sema3F (cyan) and Sema3A (magenta) expression in patient dervied tissue sections. RNA scope was combined with UEA1 fluorescent staining to identify vessels. The expression in VM lesional vessels was compared to non-lesional vessels. Scale bar: 50µm. (g) Spatial gene expression levels were quantified and expressed as % of positive signal (coverage)/nucleus. n≥100 EC nuclei/group. Mean±SD, Welch’s t-test. P-values are indicated. (h) qPCR for Sema3F and Sema3A in control EC (HUVEC and ECFC) and four VM patient-derived EC (TIE2 p.L914F mutation). Mean, quartiles, Welch’s t-test. P-values are indicated.

Furthermore, we used RNAscope^TM^ technology that allowed for the visualization of gene expression of Sema3F and Sema3A within VM patient derived tissue sections while preserving the spatial organization. Ulex europaeus agglutinin I (UEAI) fluorescent staining was used to identify lesional and non-lesional vessels within tissue sections resected from VM patients harboring a TIE2-L914F mutation (Suppl. Fig. 5). Using this system, we compared expression levels within ECs lining the different types of vessels and observed significantly (p<0.0001) higher expression levels of Sema3F and Sema3A in EC lining lesional compared to non-lesional vessels (Fig. 3f-g). Sema3A and Sema3F overexpression was furthermore confirmed in primary EC expressing the TIE2-L914F mutation which were isolated from VM patients (Fig. 3h). Therefore, we hypothesized that the overexpression of chemorepellent Sema3A and/or Sema3F observed in both patient-derived VM EC and the engineered TIE2-L914F EC can drive their competitive advantage over wild-type EC.

Moreover, wild-type EC confronted with VM patient-derived EC (with TIE2-L914F mutation ^15^) showed a migratory pattern similar to wild-type EC confronted with the engineered TIE2-L914F EC. In this setup, wild-type EC showed a reduction in FMI and directionality when in confrontation with EC from 2 out of 3 VM patients (Suppl. Fig. 6). Interestingly, the ‘repulsive’ migratory response of wild-type EC was recorded when cells were confronted with VM patient samples with higher Sema3A and Sema3F expression levels (Suppl. Fig. 6c).

### Knock-down of Sema3A or Sema3F in TIE2-mutant EC restores their repulsive effect towards wild-type EC

To evaluate the involvement of Sema3A or Sema3F in the competitive advantage, we performed shRNA-mediated knock-down of *Sema3A* or *Sema3F* in TIE2-L914F mutant EC. Knock-down was confirmed by qPCR and revealed no downregulation of Sema3A expression in Sema3F knock-down cells and vice versa (Suppl. Fig. 7). Time-lapse analysis of the cell confrontation assay revealed that the knock-down of *Sema3A* or *Sema3F* in TIE2-mutant EC revoked the repulsive effect on wild-type EC (Fig. 4a-c). Wild-type EC in the confrontation assay with TIE2-mutant EC (control vector) changed direction of migration within the confrontation phase (10-15 h), as observed previously. This confirmed the repulsive phenotype of TIE2-L914F EC towards wild-type EC (Fig. 4b), which was also represented by a significant downregulation of the forward migration index and of the directionality (Fig. 4c). These repulsive effects were not observed when wild-type EC were confronted with TIE2-WT EC, confirming that the repulsion is not caused by TIE2 overexpression but rather due to TIE2 hyperactivation due to p.L914F mutation (Suppl. Fig. 8). In contrast, when in confrontation with TIE2-L914F EC knocked-down for *Sema3A* or *Sema3F*, the wild-type EC migration behavior was completely rescued and showed migration tracks similar to the control setup (wild-type EC alone) (Fig. 4b). Quantification of migration parameters confirmed that the forward migration index and directionality of cells were normalized (Fig. 4c), revealing that knock-down of *Sema3A* or *Sema3F* can revoke the repulsive activity of TIE2-mutant EC.

**Figure 4.**
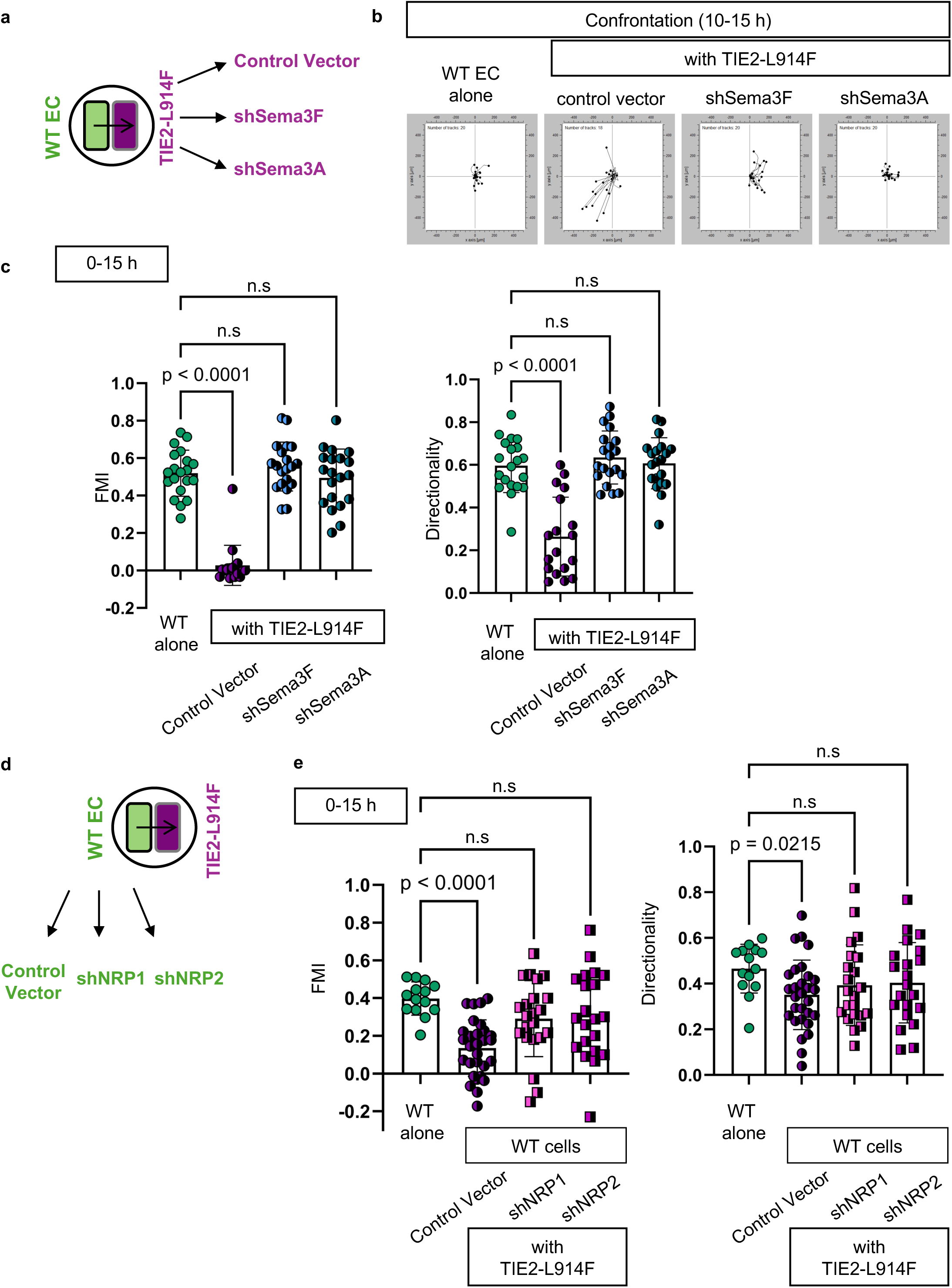
Knock-down of Sema3A/3F or NRP1/2 rescues repulsion of wild-type EC. (a) Schematic of cell confrontation assay. Wild-type EC are confronted with TIE2-L914F EC treated with control Vector or shRNA for Sema3F or Sema3A. (b) Wild-type trajectory blots in the confrontation phase (10-15h). c) Quantification of forward migration index (FMI) and directionality during the whole duration of the assay (0-15h). n≥20 cells per condition were tracked. Mean±SD, One-Way ANOVA. (d) Schematic of cell confrontation assay. Wild-type EC treated with control Vector or shRNA for NRP1 or NRP2 are confronted with TIE2-L914F EC. (e) Quantification of forward migration index (FMI) and directionality during the whole duration of the assay (0-15h). n≥14 cells per condition were tracked. Mean±SD, One-Way ANOVA. P-values are indicated.

### Knock down of NRP1 or NRP2 in wild-type EC prevents repulsive effect of TIE2-L914F cells

Sema3A and Sema3F signal through Neuropilin-1 (NRP1) or NRP2 in a complex with Plexin A receptors ^29,54–56^. Analysis of receptor expression by immunoblotting revealed NRP1 expression was slightly decreased in TIE2-L914F EC compared to wild-type EC and TIE2-WT EC, while NRP2 levels were comparable (Suppl. Fig 9a-b). Analysis of Plexin A receptors in mutant and WT EC revealed similar expression levels for *PLXNA1* and *PLXNA2* while *PLXNA3* and *PLXNA4* gene expression was downregulated in TIE2 mutant EC (Suppl. Fig. 8c).

To investigate the receptors involved in the Sema3A/3F chemorepulsion response in wild-type EC, we knocked-down *NRP1* or *NRP2* (Suppl. Fig. 10) in wild-type EC and confronted them with TIE2-L914F cells. When wild-type EC knocked-down for *NRP1* or *NRP2* were confronted with TIE2-L914F cells (Fig. 4d), their migration behavior was rescued. Quantification of migration parameters confirmed that the forward migration index and directionality of cells were normalized (Fig. 4e), supporting the hypothesis that the repulsive activity of TIE2-mutant EC is modulated through the Sema3/NRP axis.

### Knock-down Sema 3A or 3F in TIE2-L914F EC rescues the ability of wild-type EC to sprout and form lumens

To evaluate if the knock-down of *Sema3A* or *Sema3F* restores the ability of wild-type EC to sprout and form lumens, a 3D cell competition assay was setup by coating beads separately with fluorescently labeled wild-type EC or TIE2-L914F EC, treated with control vector or shRNA for *Sema3A* or *Sema3F* (Fig. 5a). At the endpoint, the 3D cultures were imaged by z-stack confocal imaging and vascular structures were reconstructed using Imaris imaging software (Fig. 5b). In the presence of TIE2-mutant EC treated with control vector, the number of wild-type EC sprouts per bead was significantly inhibited compared to wild-type EC alone (Fig. 5c). In contrast, the knock-down of *Sema3A* or *Sema3F* in TIE2-mutant EC rescued wild-type EC sprouting, partially restoring their morphogenesis and lumen formation ability (Fig. 5b,c).

**Figure 5.**
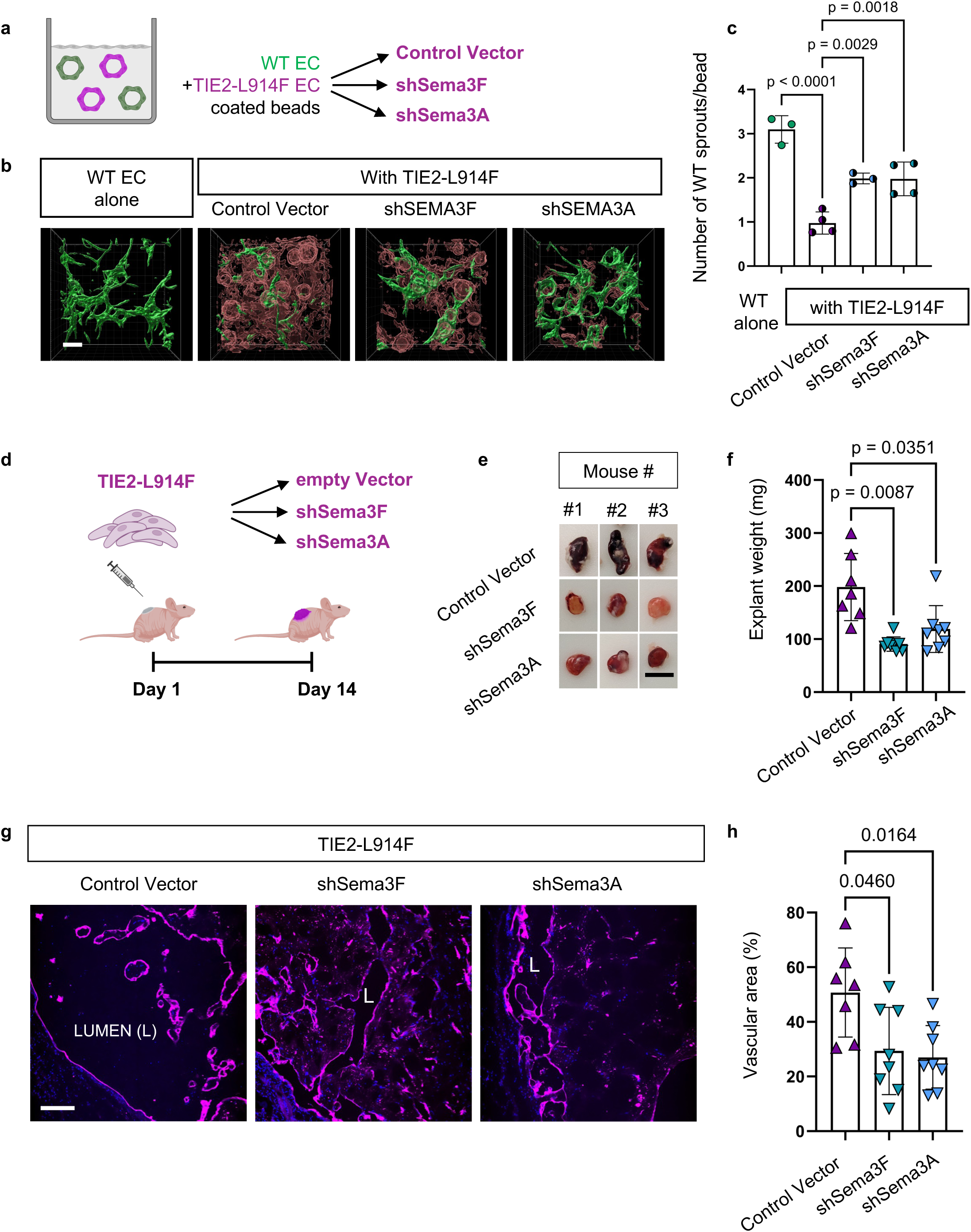
Knock-down of Sema3A or Sema3F restores wild-type EC sprouting. (a) Dextran beads are separately coated wild-type EC (shown in green), or TIE2-mutant EC (shown in magenta) treated with control Vector, shSema3A or shSema3F and combined in the 3D cell competition assay. (b) 3D reconstruction of confocal z-stack images on day 8. Scale bar: 200µm. (c) Quantification of number of HUVEC sprouts per bead. Mean±SD, One-way ANOVA. (d) Schematic of VM xenograft model with TIE2-L914F EC treated with control Vector or shRNA for Sema3A or Sema3F. In this model TIE2-L914F EC are injected alone to assess the cell autonomous effects of Sema3A and 3F. (e) Representative photographs of the VM xenograft explants at day 14. Scale bar: 1cm (f) Weight of xenograft explants. n≥7 mice/condition. Mean±SD, Welch’s t-test. (g) Xenograft tissue sections stained for UEAI (Ulex europaeus I, magenta). Lumen (L) indicated. Scale bar: 200µm. (h) Quantification of vascular area (%). n≥7 mice/condition, mean±SD, One-way ANOVA. P-values are indicated. Schematic in (a and d) created with Biorender.com.

### Sema3A and Sema3F exhibit cell autonomous effects on TIE2-L914F EC

Next, we explored the cell autonomous effects of Sema3F and Sema3A in TIE2-mutant EC in the VM xenograft model. Fluorescently labeled TIE2-L914F EC (control vector, shSema3F or shSema3A) cells were suspended within a Matrigel matrix and injected alone into the flanks of nude mice (Fig. 5d). Xenograft explants were analyzed at day 14 (Fig. 5e). Macroscopically, the explants containing TIE2-L914F EC (control vector) were visibly perfused indicating pathological vessel formation and growth. In contrast, the explants of TIE2-L914F EC knocked-down for *Sema3A* or *Sema3F* were visibly less vascularized and significantly (p<0.05) lighter in weight (Fig. 5e-f, Suppl. Fig. 11a-b), indicating an inhibition of vessel growth. Histologically, TIE2-mutant cells, knocked-down for *Sema3F* or *Sema3A,* formed xenografts with a significantly (p<0.05) reduced vascular area (Fig. 5g-h) compared to control vector. Proliferation analysis of TIE2-L914F EC knocked-down for either *Sema3A* or *Sema3F* showed there was no significant difference in the proliferation rates, suggesting that the reduction of lesion size is not due to differences in cell proliferation (Suppl. Fig. 11c). These results indicate that Sema3A and Sema3F also play a cell autonomous role in TIE2-mutant EC during the lumen enlargement.

### Knock-down of Sema 3A or 3F in TIE2-L914F EC restores cell communication with wild-type EC and reduces vascular lesion size

Our data suggest that VM lumens are formed by clonal expansion of TIE2-mutant EC and show that wild-type EC are repulsed and excluded from the aberrant VM vascular lumens. While downregulation of *Sema3A* or *3F* in TIE2-L914F EC restored migration and sprouting of wild-type EC, it is paramount to further determine if: 1-recruitment of wild-type EC to the mutant blood vessels could rescue the abnormal VM vessel morphology or 2-could instead be detrimental by promoting further lumen enlargement (Fig. 6a).

**Figure 6.**
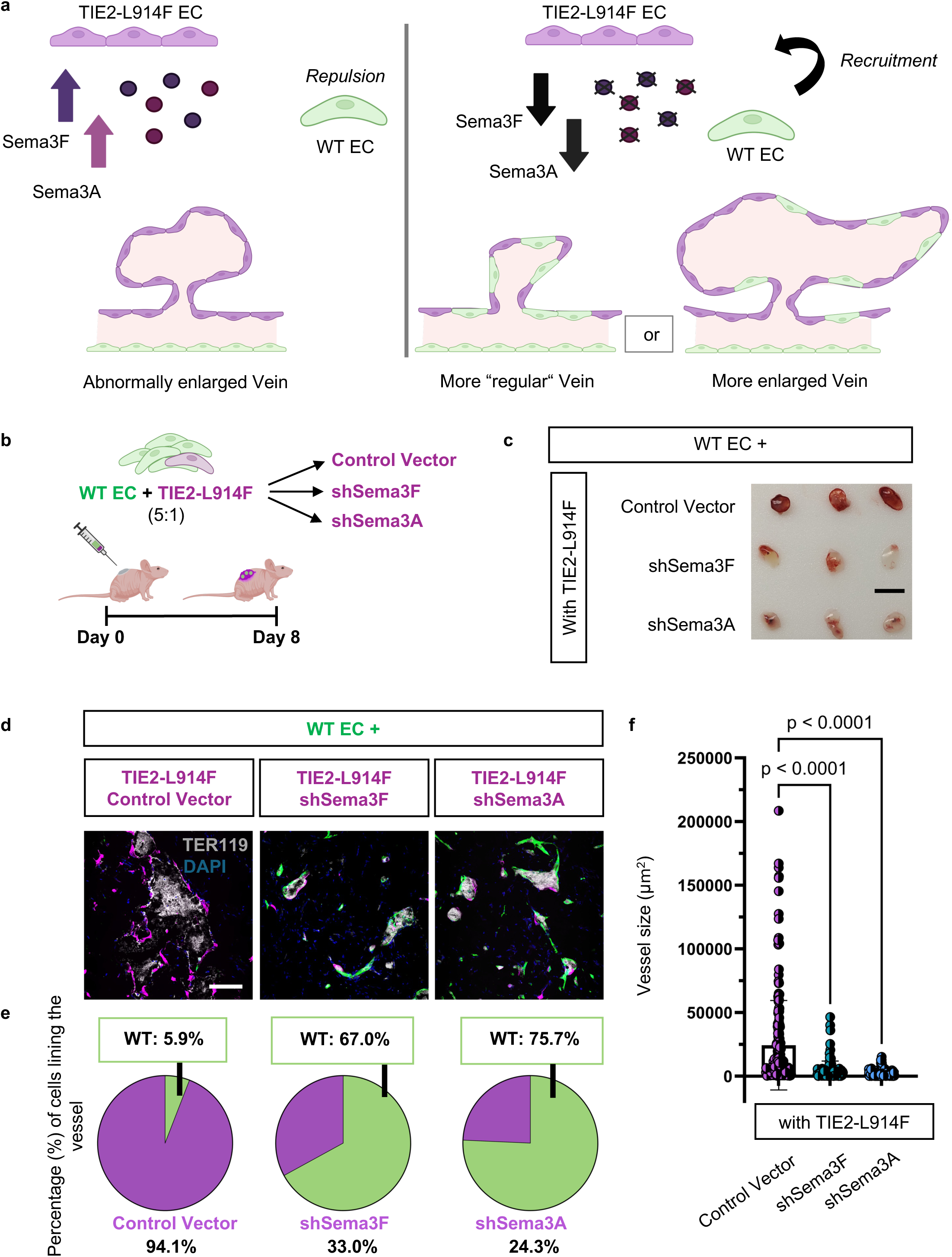
Knock-down of Sema3A or Sema3F restores in vivo communication between TIE2-mutant EC and wild-type EC, reducing VM vascular lesion size. (a) Schematic of possible scenarios. TIE2-L914F cells express high levels of chemorepulsive molecules Sema3F and Sema3A that are responsible for the repulsion of wild-type EC (left panel). We hypothesize that if we inhibit Sema3A or Sema3F (right panel) wild-type cells will no longer be repulsed and incorporated into the lesion vessel. This recruitment of wild-type EC could lead to normalization of the vessel size or promote further lumen enlargement. (b) Schematic of the mixed EC xenograft model of VM. Wild-type EC intermixed with HUVEC TIE2-L914F (control Vector, shSema3A, or shSema3F), dissected on day 8. (c) Representative photograph of xenograft explants. Scale bar: 1cm. (d) Explant tissue sections with wild-type EC shown in green. TIE2-L914F EC treated with either control Vector, shSema3F or shSema3A are represented in magenta. TER119 shows erythrocytes (grey) confirming perfusion of vessels. Scale bar: 200µm (e) Quantification of the cell type lining vessels and (f) Quantification of vessel size. n=8 mice/condition and n≥115 vessels/condition, mean±SD, One-way ANOVA. P-values are indicated. Schematic in (a,b) created with Biorender.com.

To address this question, we performed xenograft studies to determine the lumen size effects of restoring wild-type/mutant EC crosstalk in vivo. We intermixed fluorescently labeled wild-type EC (shown in green) with TIE2-L914F EC (shown in magenta) knocked-down for *Sema3F* or *Sema3A* and injected them into mice (Fig.6b). Explants containing TIE2-mutant EC knocked down for *Sema3A* or *Sema3F* were visibly smaller and less perfused compared to xenograft explants of the control vector group (wild-type EC mixed with TIE2-L914F control vector) (Fig. 6c, Suppl. Fig. 12a-b). The weight of xenograft explants was significantly (p<0.05) reduced compared to lesions containing TIE2-L914F EC treated with control vector (Suppl. Fig. 12c). Next, we analyzed the % of wild-type EC in the VM vascular lumens. Blood vessels in the control setup were lined almost exclusively by TIE2-mutant EC (control vector) (94.1%), and only a few wild-type EC lined the ectatic VM-like lumens (5.9%) (Fig. 6d-e, Suppl. Fig. 12d). When *Sema3F* or *Sema3A* were knocked-down, the interaction between TIE2-L914F and wild-type EC was restored, and EC lining blood vessels were comprised of 67 or 75.7% of wild-type EC, respectively (Fig. 6d-e, Suppl. Fig. 12d). Importantly, vessel size analysis demonstrated that the recruitment of wild-type EC into the VM vessels was accompanied by reduced vessel size in the xenograft plugs (Fig. 6f). Analysis of wild-type EC recruitment in small (≤ 100μm) versus large (> 100μm) vessels in the control group did not reveal significant difference (Suppl. Fig. 12e). This indicates that the increased recruitment of wild-type EC is not related to the reduced vessel size but rather to the knockdown of *Sema3A* and *Sema3F*. These results suggest that recruitment of wild-type EC can normalize the ectatic VM vessel morphology, while repulsion of wild-type EC is a mechanism implicated in the pathogenic VM lumen expansion. Altogether, our data show that Sema3A and Sema3F play an important role in VM pathogenesis and we propose here that their inhibition can reduce lesion burden.

## Discussion

To date, little research has focused on the cellular and molecular mechanisms involved in the abnormally enlarged blood vessel formation involved in the pathogenesis of TIE2-mutated VM. Here, using a VM xenograft model, we showed that lesional vessels were almost exclusively lined by TIE2-mutant EC while wild-type EC were mostly absent. With a 3D cell competition assay and 2D cell confrontation assay, we have shown that TIE2-mutant EC inhibited the ability of wild-type EC to sprout and migrate while promoting their cell repulsion. This effect was mediated by overexpression of Sema3A and Sema3F. Knock-down of these chemorepellent molecules restored communication of TIE2-mutant EC with wild-type EC and rescued the VM phenotype by suppressing vascular lumen enlargement.

The key question of this study was whether the aberrant venous expansion is caused by the recruitment of wild-type EC into the growing VM lesion. A similar question has been raised by other investigators while studying the development of cerebral cavernous malformations (CCM). In CCM, brain lesions are characterized by cavernous lesions with ectatic blood vessels caused by loss-of-function mutations in any one of three CCM genes ^57^. Both VM and CCM pathogenesis are characterized by massively enlarged venous channels. In addition, in CCM, as in other vascular malformations including VM, causative somatic mutations are commonly found at low (<10%) allelic frequency in patients’ affected tissue ^58^. This low mutant allelic frequency has prompted investigations into the cellular contributions to the abnormally enlarged blood vessels. Recently, two independent, groundbreaking studies have elegantly shown, with the use of a Confetti reporter, that CCM mutant cells in transgenic mice form enlarged blood vessels by initial clonal expansion followed by recruitment of wild-type EC into the growing lesion ^59,60^. The recruitment of wild-type EC into CCM lesions was further demonstrated by mosaicism of CCM-null (mutant) and wild-type EC in large lesions from CCM patients ^61^.

In contrast to these findings in CCM, our study demonstrates that in TIE2-mutant VM the pathological expansion of the veins is not defined by recruitment of wild-type EC. Instead, TIE2-L914F EC can promote repulsion and inhibition of morphogenesis in wild-type EC (competitive advantage) to propel their own clonal expansion. In addition, the reduced recruitment of wild-type EC was independent of vessel size as no difference was detected in small versus large vessels.

The contribution of EC proliferation (i.e. clonal expansion) to the VM phenotype is still debated. Historically, when compared to vascular tumors, EC proliferation in VM has not been considered pathological ^50,62^. For this reason, our initial hypothesis was that recruitment of wild-type EC would be a mechanism allowing pathological VM lumen expansion. Conversely, our results here show that clonal expansion has an important role in VM pathogenesis and can regulate the progressive nature of VM lesions. This is in line with recent studies highlighting the hyperproliferative nature of VM blood vessels ^13,63,64^. As VMs are characterized by abnormally enlarged vascular channels and are active lesions that grow and recur, it is reasonable to assume that increased EC proliferation has an important role. Here, we would like to speculate that proliferation may be overlooked in clinical VM biopsies as lesions are only resected when they reach a certain size that may not reflect all stages of VM initiation and progression.

In seeking to define the mechanism driving the competitive advantage, we detected elevated expression levels of chemorepellent Sema3A and Sema3F in the TIE2-mutant EC. While the role of Semaphorins in VM pathogenesis has not been investigated prior to this study, the upregulation of Sema3A and Sema3F expression in TIE2-L914F mutant EC has been reported in previously published microarray datasets (GSE46684) ^17^, hence confirming our findings.

Sema3A has been shown to have an important role in the patterning of veins ^65^. This may be related to its ability to inhibit EC migration in vitro and sprout formation that we show here and has also been reported during retinal vasculature development ^30^ ^20,23^. Sema3F functions as an inhibitor of tumor angiogenesis ^22,66^ by causing cytoskeleton collapse ^67^ and inhibition of EC survival ^66^. While the anti-angiogenic effects of Sema3A and 3F have been linked with EC survival inhibition ^23^, in our study wild-type EC did not seem to undergo cell death as they persisted in the 3D fibrin gel cell competition assay until day 14 and treatment of wild-type EC with TIE2-L914F conditioned media did not significantly change wild-type cell number. In our xenograft model, we demonstrated that the wild-type EC do not incorporate into a functional blood vessel in presence of TIE2-mutant EC. Matrigel-based xenografts generated with wild-type EC do not show vessel formation and EC do not persist in the xenografts when not incorporated into a functional vessel ^68^. Thereby, in our xenograft system, it is not possible to conclude whether wild-type EC survival is negatively impacted by exposure to Semaphorin from mutant EC or merely by exclusion from functional vessels.

Our in vitro data show a non-cell autonomous role for Sema3A and Sema3F in VM. ShRNA-mediated knock-down of Sema3A or Sema3F in TIE2-mutant EC rescued the repulsive migration phenotype of wild-type EC and reversed the inhibitory effect towards wild-type EC sprouting in a 3D cell competition assay. These results were confirmed in a murine VM xenograft model where knock-down of Sema3A or Sema3F promoted recruitment of wild-type EC into the vessels and significantly reduced the vessel size. In published in vitro studies, the repulsive and anti-proliferative effects of Sema3F have been shown to be synergistically enhanced by Sema3A ^23^. Interestingly, in our study, the single knockdown of Sema3A or Sema3F was sufficient to completely rescue the repulsive migratory phenotype. This suggests that the combination of both Sema3A and 3F signaling is not necessary for the full repulsive effect.

Histologically, VM enlarged channels are characterized by abnormal and discontinuous perivascular/smooth muscle cell coverage. In the in vivo environment, the lack of perivascular coverage in a vein could allow for the hydrostatic pressure to promote vessel thinning and enlargement. Decreased levels of platelet derived growth factor-B (PDGF-B) in TIE2-mutant EC have been linked to decreased pericyte/smooth muscle cell recruitment ^17^. However, other mechanisms could contribute to this phenotype, and it is interesting to speculate that the increased levels of Sema3A and 3F released by the TIE2-mutant EC could play a role in the inhibition of pericyte recruitment (or in promoting pericyte repulsion) in the pathological VM veins. This hypothesis, in conjunction with our results, suggests a potential role for therapies targeting Sema3A or Sema3F in the management of VM. In the context of vascular diseases, Sema3A inhibition has been shown to be a promising strategy for the therapeutic treatment of diabetic retinopathy and macular ischemia, retinal vein occlusion, and macular edema in AMD ^69–71^. Further studies to assess the preclinical efficacy of Sema3A or 3F inhibition strategies in VM models are warranted.

In closing, we would like to discuss a few limitations of our study. 1-While we demonstrated the importance of clonal expansion and Sema3A/3F during the enlargement of TIE2-L914F-related VM lesions, it is still to be determined if these mechanisms are translatable to other *TIE2* or *PIK3CA* mutations associated with VM. Interestingly, mutant *PIK3CA*-based genetic murine models of VM showed high proliferation in the VM-like vascular lesions in the early postnatal retina, partially supporting our findings ^13,64,72^. Sema3F is negatively regulated by the transcriptional regulator Forkhead box O 1 (FOXO1)^73^. Both hyperactive TIE2 or PIK3CA mutations result in increased AKT activation which, in turn, leads to FOXO1 phosphorylation^74^ and cytoplasmic sequestration. This knowledge suggests that PIK3CA mutant EC could also upregulate Sema3F, which can be a common pathogenic mechanism in both TIE2 and PIK3CA mutant VM. Furthermore, treatment with the mTOR inhibitor rapamycin or the PIK3CA inhibitor Alpelisib have been shown to suppress AKT activation in VM EC^12,51,75^, this would suggest that their partial efficacy in patients could include a reduction in Sema3F expression. 2-Here, we have used a combination of 2D, 3D and xenograft strategies to demonstrate the competitive advantage TIE2-L914F EC over wild-type EC. Future studies should focus on the generation of transgenic / knock-in mutant TIE2 murine models which can more faithfully recapitulate the developmental stages of the VM pathogenesis. These models would allow for the tracking of mutant and wild-type cells at different stages of vessel formation and enlargement to implicate more definitely the clonal expansion (and lack of mosaicism) of TIE2-mutant EC in VM disease.

## Acknowledgements and Sources of Funding

Research reported in this manuscript was supported by the National Heart, Lung, and Blood Institute, part of the National Institutes of Health, under Award Number 2R01 HL117952 (E.B.), R01 HL167700 (E.B.) and F31 HL176101 (L.B.). The content is solely the responsibility of the authors and does not necessarily represent the official views of the National Institutes of Health. Additional funding supporting the study was provided by the American Heart Institute (AHA) Postdoctoral Fellowship (award number 833891) to S. Schrenk. We thank Rachael Kang for technical assistance with cryosections. We thank the Bio-imaging and Analysis Facility, Bioinformatic Core and Veterinary Services at Cincinnati Children’s Hospital Medical Center and the Genomics, Epigenomics and Sequencing Core at the University of Cincinnati for providing state-of-the-art instrumentation, services, training, and education.

## Author contributions

S.S and E.B. designed the study. S.S. performed the experiments and data analysis. Y.C., C.S and L.B performed experiments. S.S and E.B wrote the manuscript. All authors revised and commented on the manuscript.

