## Supplemental data for "Semaphorin 3A and 3F overexpression in TIE2 hyperactive endothelial cells contribute to the pathological lumen expansion in venous malformation"

**Content:**

Supplementary Table 1

Supplemental Figure 1-12 and Supplementary Figure legends

Other Supplementary Materials for this manuscript include the following:

Supplementary Movies 1-2

**Supplementary Video 1. Cell migration of wild-type EC alone**

The migration of wild-type EC toward wild-type EC was visualized through time-lapse imaging in 30min intervals for a total of 15h. Cell nuclei were visualized and shown in white. Individual cells (≥ 20 cells/condition) were tracked over time using manual cell tracking. Each individual cell is represented by a different color

**Supplementary Video 2. Cell migration of wild-type cells towards TIE2 mutant EC**

The migration of wild-type EC toward TIE2-L914F EC was visualized through time-lapse imaging in 30min intervals for a total of 15h. Cell nuclei were visualized and shown in white. Individual cells (≥ 20 cells/condition) were tracked over time using manual cell tracking. Each individual cell is represented by a different color

**Supplementary Table 1. Real-time PCR Primers**

| | Forward (5'-3') | Reverse (5'-3') |
|---|---|---|
| **SEMA3A** | TCC CAC TGC AAA GAG ACG CA | GCT CTC TGC GAC TTC GGA CT |
| **SEMA3F** | GAT GCA GCC ATC TTC CGC AC | AGC ATG GAT GAA CGA CGG GT |
| **HPRT1** | CCT GGC GTC GTG ATT AGT GAT | GGG CTA CAA TGT GAT GGC CT |
| **TBP1** | GTG GGG AGC TGT GAT GTG AA | TGC TCT GAC TTT AGC ACC TGT |
| **NRP1** | ACA CCT GAG CTG CGG ACT TT | GGC CTG GTC GTC ATC ACA C |
| **NRP2** | CGG CTT TTG CAG GTG AGA ATT T | TCC AGT CCA CCT CGT ATT CAT CAT C |
| **PLEXA1** | CAT TGA GAG GGA GAA CGG CT | CAC GTT GTC CAT GAC GAA GC |
| **PLEXA2** | GAT CCG GAG AGA GAC CCC GA | TTT TCC AGCGCG ACT TTC CG |
| **PLEXA3** | CGG AGT ACT TCC CCA CCT TG | ACT CAT CCT GGT ACA CGA GAC |
| **PLEXA4** | AGC GTC CAC AAT TCG TGC CT | ACG GGC ACC AGG ATC TTG TC |

### Supplementary figures

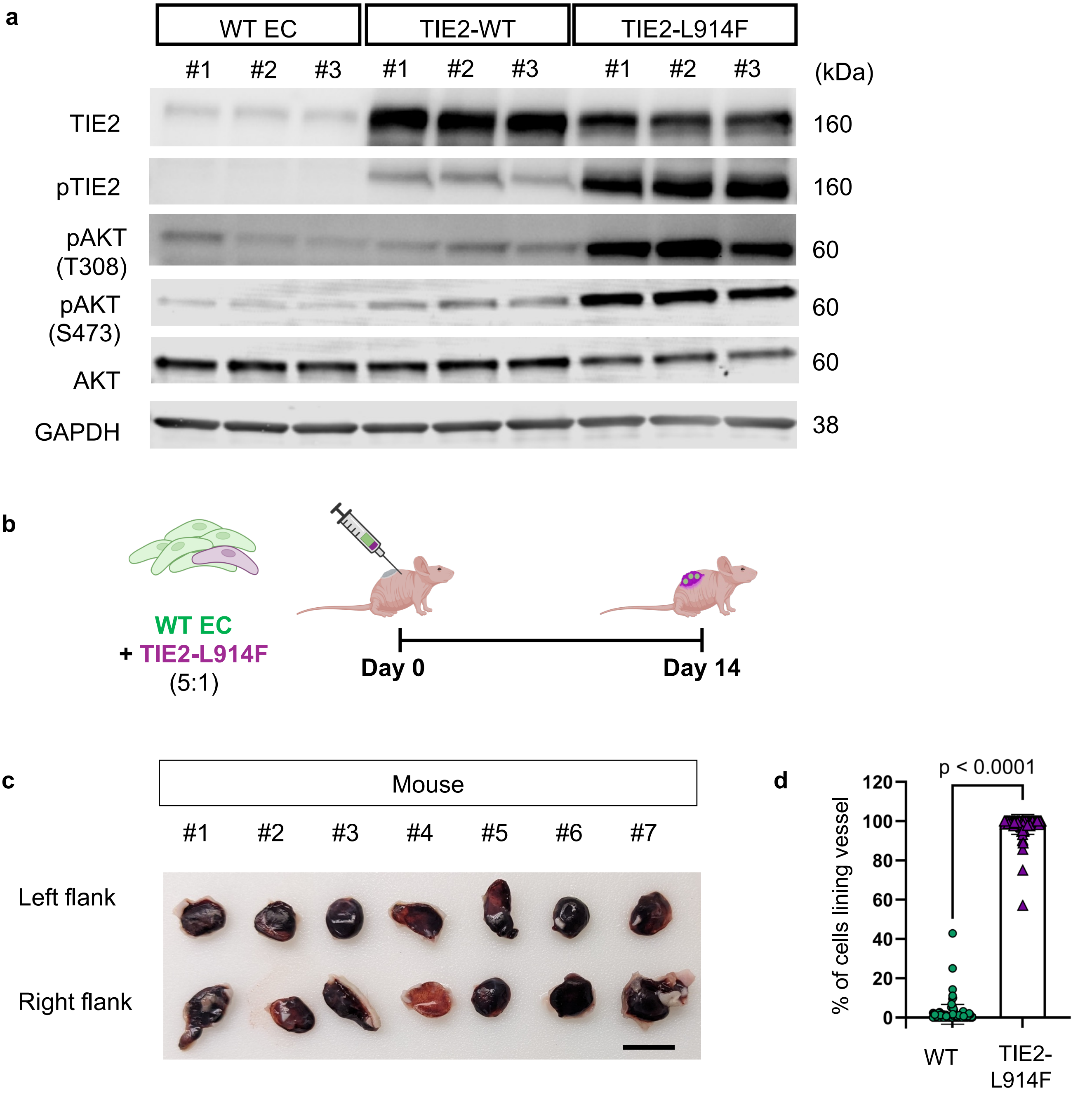


**Supplemental Figure 1. In vitro model and mixed EC xenograft model of VM.** (a) Immunoblot of wild-type (WT) EC (HUVEC), TIE2-WT (HUVEC) and TIE2-L914F EC (HUVEC) of the following targets: TIE2, pTIE2, pAKT (T308 and S473), AKT and GAPH as a loading control. n=3 biological replicates. (b) Schematic of the mixed EC xenograft model of VM: wild-type EC (green) were intermixed with TIE2-L914F EC (magenta) at a 5:1 ratio and injected in the flanks of immunocompromised nude mice. Schematic created with Biorender.com (c) Photograph of xenograft explants at day 14 after cell injection. n=7mice. Scale bar: 1cm. (d) Quantification of the cell types lining the ectatic blood vessels shown as % of cells lining vessel. n=7 mice and n=122 vessels. Mean±SD, Welch's t-test. P-values are indicated.

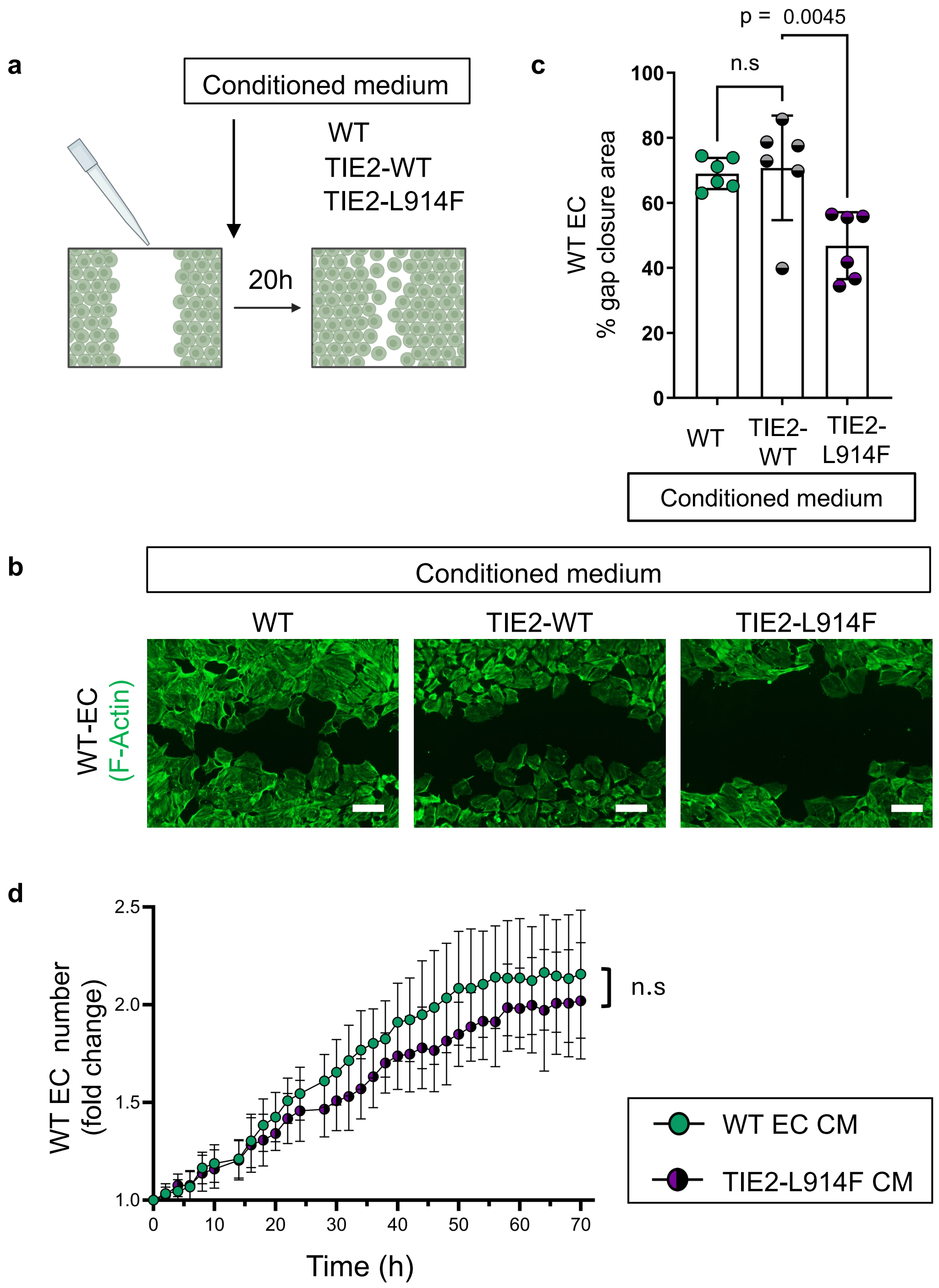


**Supplemental Figure 2. Migration of wild-type EC is inhibited by conditioned medium from TIE2 mutant EC.** (a) Schematic of the scratch wound healing assay. A scratch was produced in a confluent cell monolayer of wild-type EC, and they were exposed to conditioned medium collected form monolayers of WT EC, TIE2-WT EC or TIE2-L914F EC. Schematic was created with Biorender.com. (b) Cells were fixed 20h after the scratch was made and conditioned medium was added. F-Actin (green) is visualized. Scale bar: 200µM (c) Quantification of WT EC gap closure. n=6 independent experiments. Mean±SD, One-way ANOVA. p-values are indicated. (d) Effect of conditioned medium on wild-type EC proliferation was analyzed via time-lapse imaging over a time course of 70h. n=8 wells per condition. Mean±SD, 2-way ANOVA.

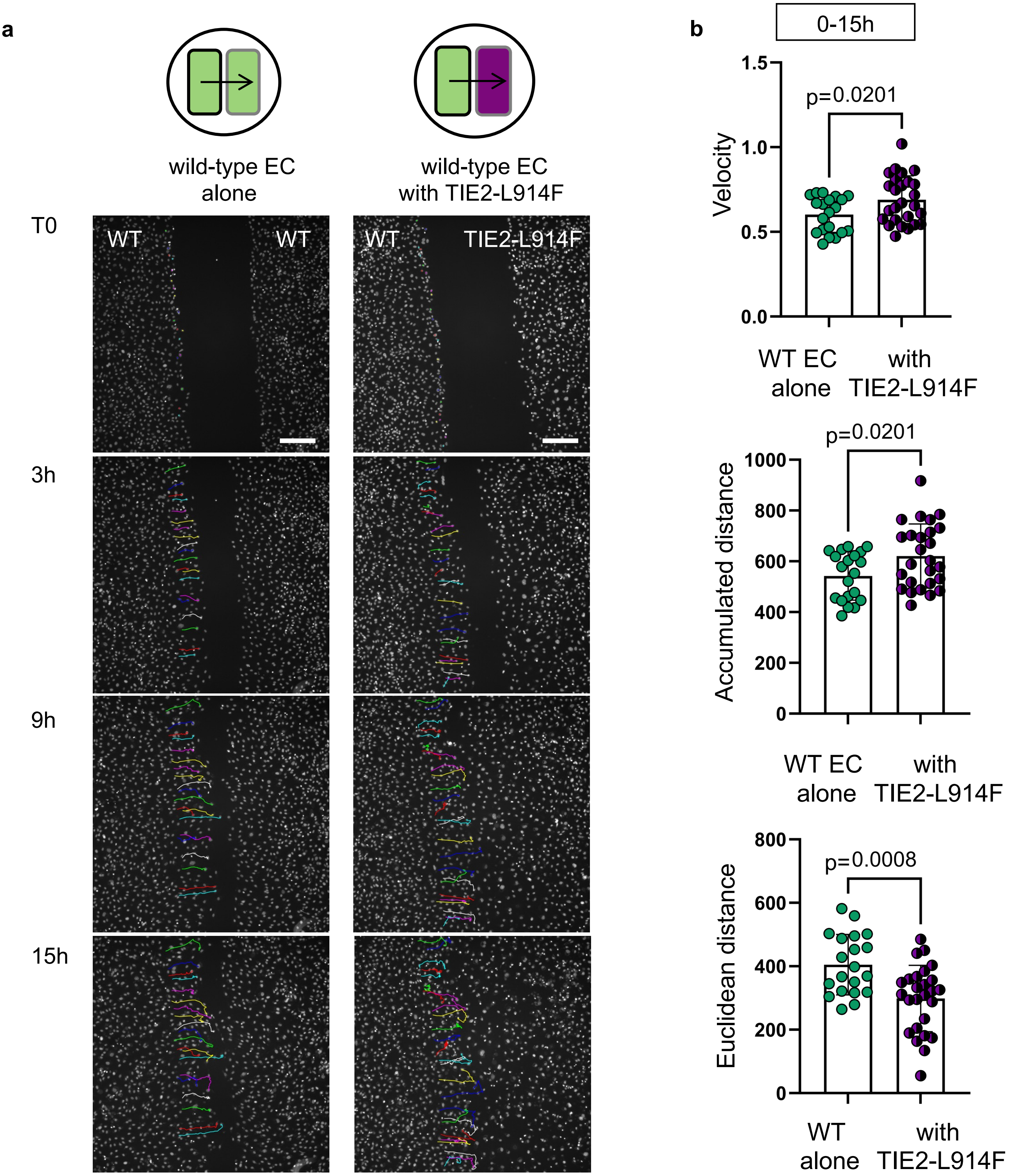


**Supplemental Figure 3. TIE2 mutant EC repulse wild-type EC.** (a) Schematic of cell confrontation assay. The migration of wild-type EC toward WT EC (left panel) or TIE2 L914F EC (right panel) was visualized through time-lapse imaging and then analyzed via cell tracking. Each tracked cell is represented by a different color at T0, 3h, 9h and 15h. Scale bar: 200µM. (b) Quantification of velocity, accumulated distance and euclidean distance. n≥20 cells per condition were tracked. Mean±SD, Welch's t-test. p-values are indicated.

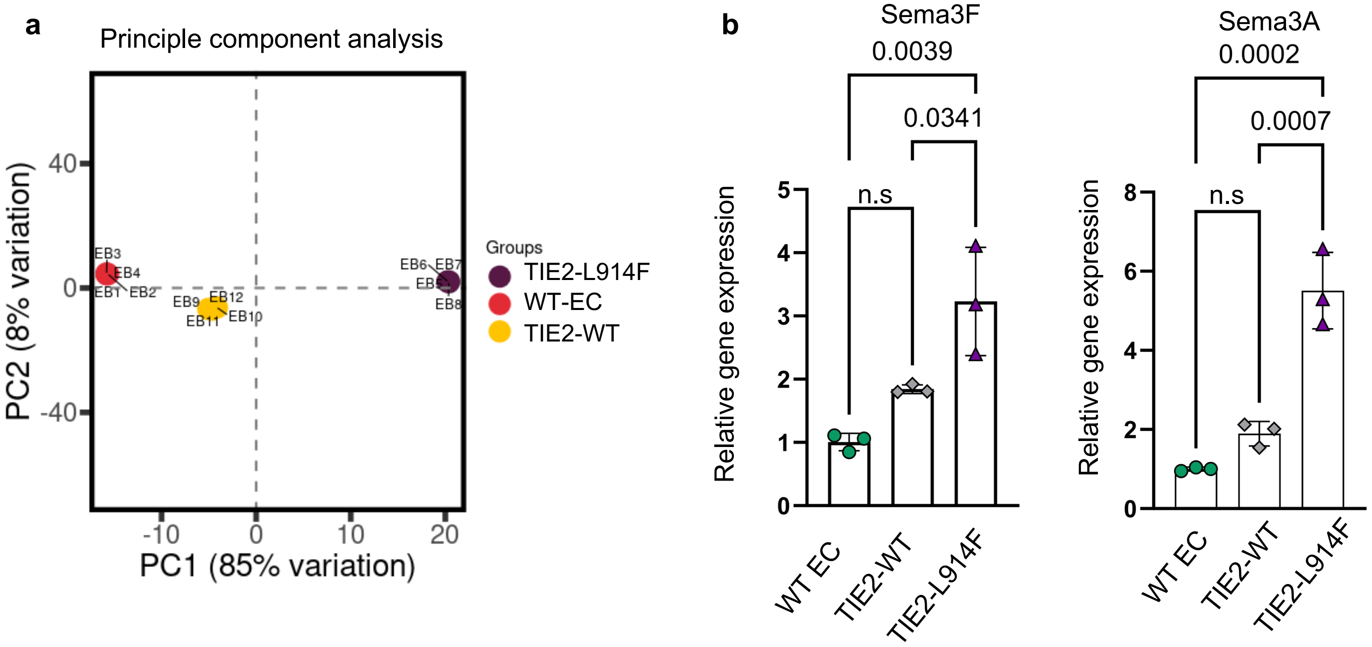


**Supplemental Figure 4. RNA-Sequencing of TIE2 L914F EC reveals enriched expression of genes related to chemorepulsive semaphorin expression**. (a) Principal Component Analysis (PCA) of RNA-sequencing data obtained from WT EC (red), TIE2-WT EC (yellow) and TIE2-L914F EC (magenta) (n = 4 biological replicates per group). (b) Sema3F and Sema3A RNA expression levels were quantified by qPCR. n=3 biological replicates to confirm RNA sequencing results. Mean±SD, One-way ANOVA. p-values are indicated.

a

b

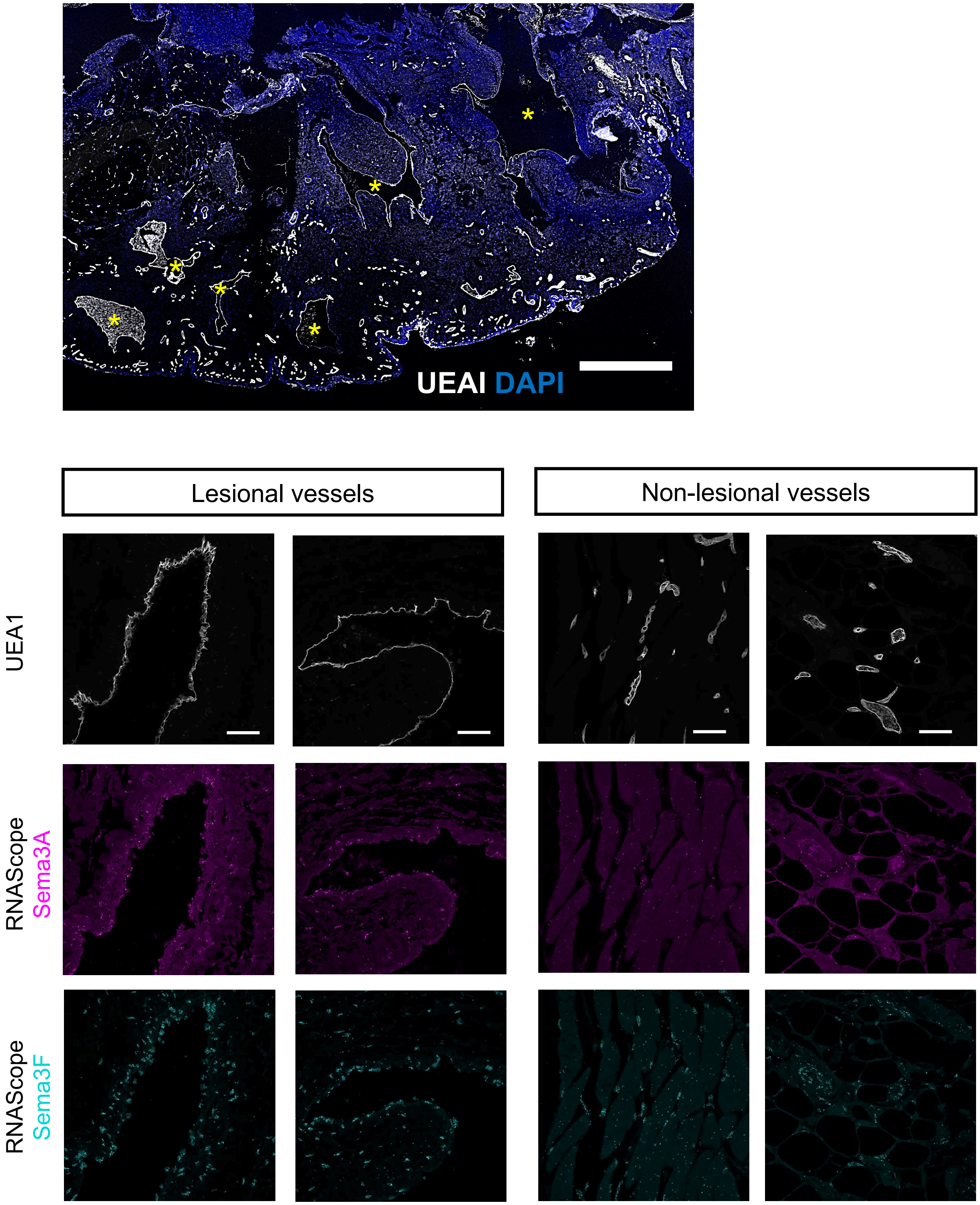


**Supplemental Figure 5. Histology of resected tissue from** VM patient used for RNA scope analysis. (a) Overview fluorescent staining using UEAI (white and DAPI) shows irregular, abnormally enlarged lesional vessels (yellow star) as well as small, non-lesional vessels within resected tissue. Scale bar: 1cm. (b) Higher magnification representative images of RNA scope for Sema3A and Sema3F combined with UEA1 fluorescent staining (white) in lesional and non-lesional vessels. Scale bar: 50 μm

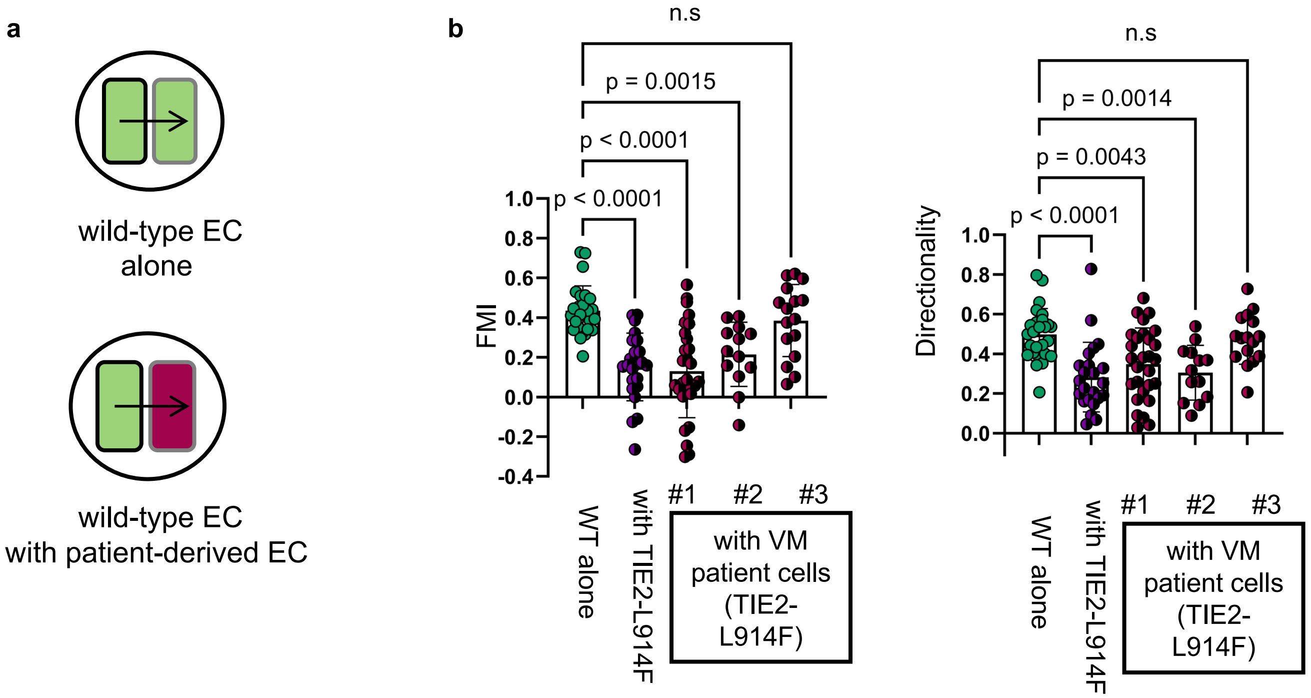


**c**

| | VM#1 | VM#2 | VM#3 |
|---|---|---|---|
| Sema3A gene expression fold change | 3.22 | 3.39 | 1.13 |
| Sema3F gene expression fold change | 2.93 | 2.92 | 1.30 |

**Supplemental Figure 6. VM patient-derived EC with increased Sema3A and Sema3AF levels repulse wild-type EC.** (a) Schematic of cell confrontation assay. The migration of wild-type EC toward WT EC (upper panel) or VM patient derived EC (TIE2 L914F mutation) (lower panel) was visualized through time-lapse imaging and then analyzed via cell tracking. (b) Quantification of forward migration index (FMI) and directionality over the timecourse of 15h. n≥20 cells per condition were tracked. Mean±SD, Welch's t-test. p-values are indicated. (c) Relative gene expression levels in VM derived patient EC expressed as fold change compared to control EC (HUVEC).

**a**

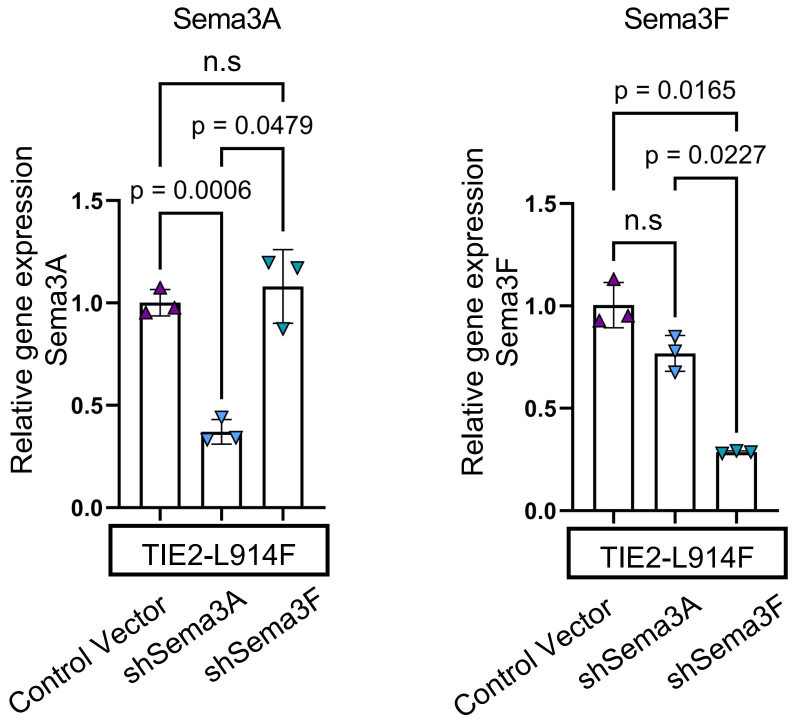


**Supplemental Figure 7. sh-RNA mediated knockdown of Sema3F and Sema3A.** (a) Quantification of Sema3A and Sema3F gene expression levels in TIE2-L914F EC treated with either control Vector, shSema3A or shSema3F Vector by qPCR. Mean±SD, One-way ANOVA. p-values are indicated.

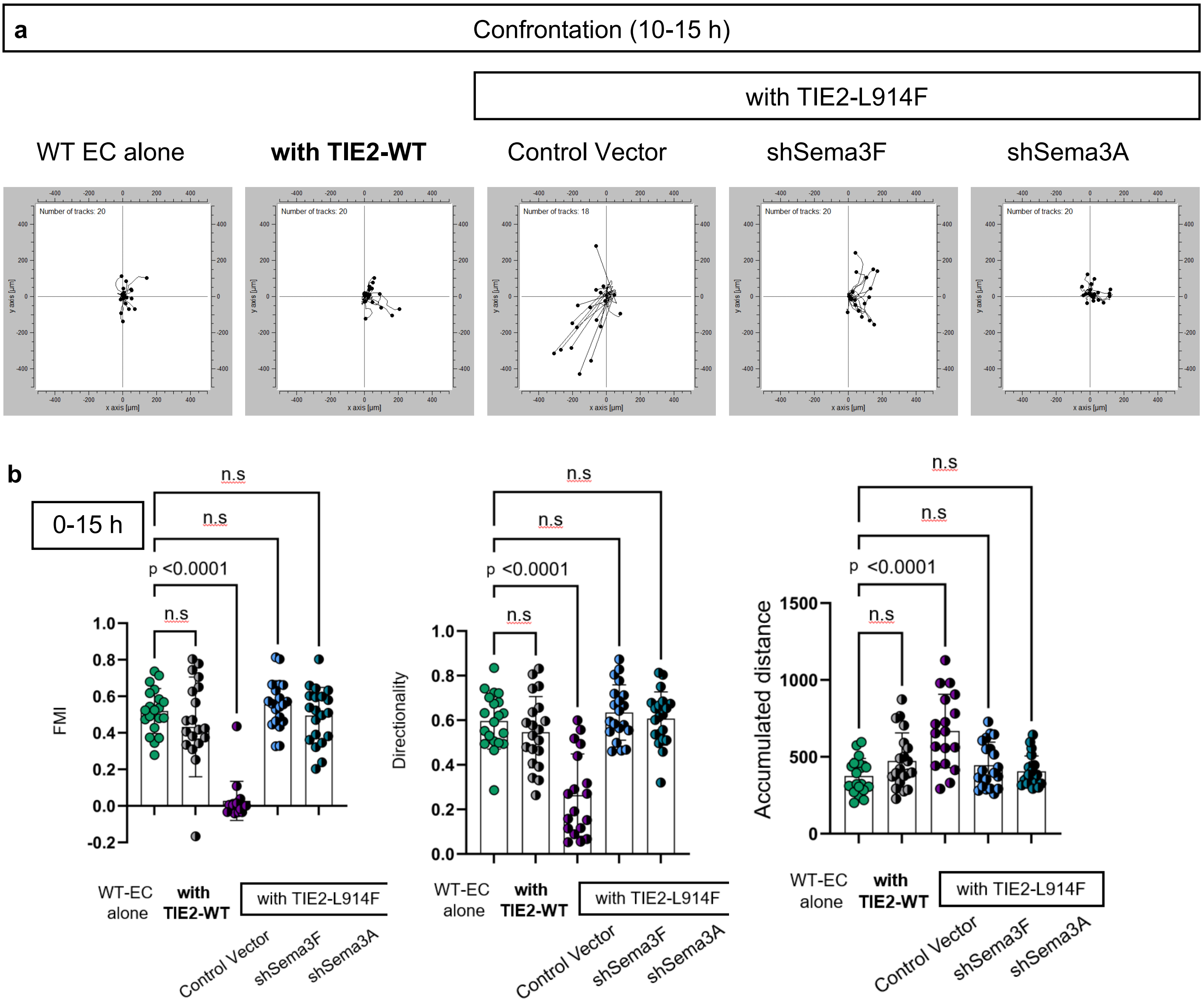


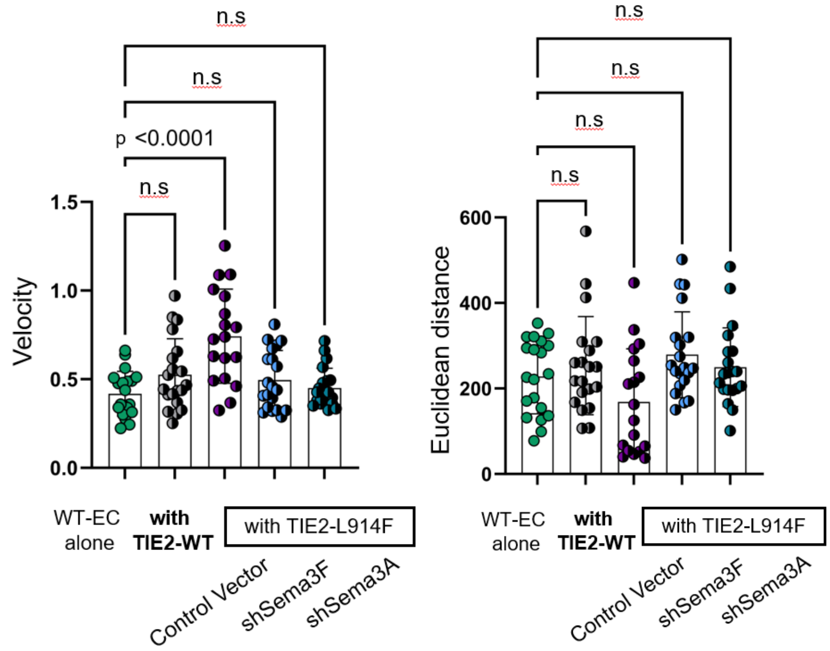


**Supplemental Figure 8. TIE2 mutant EC repulse wild-type EC.** (a) Wild-type EC are confronted with TIE2-WT EC or with TIE2-L914F EC treated with control Vector or shRNA for Sema3F or Sema3A. Wild-type trajectory blots in the pre-collision phase (10-15h). (b) Quantification of forward migration index and directionality, accumulated distance, euclidean distance, and velocity (0-15h). n≥20 cells per condition were tracked. Mean±SD, One-Way ANOVA. P-values are indicated.

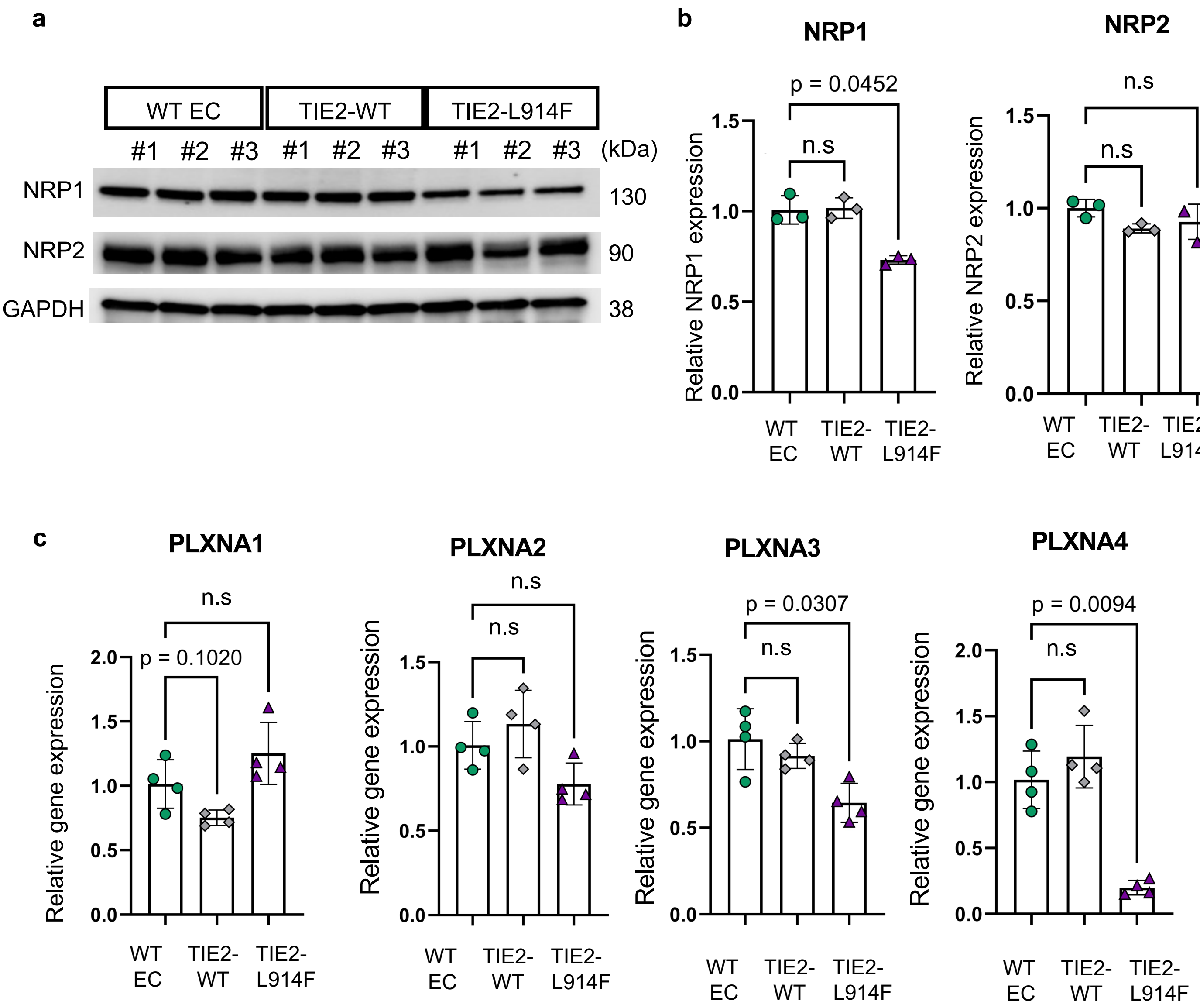


**Supplemental Figure 9. Expression levels of Sema3A and 3F receptors.** (a) Immunoblot for NRP1 and NRP2 expression in WT EC, TIE2-WT EC and TIE2-L914F cells and (b) quantification of protein expression. N=3 biological replicates. Mean±SD. One-way ANOVA. p-values are indicated. (c) Gene expression levels of NRP1, NRP2, Plexin A, Plexin A2, Plexin A3 and Plexin A4 were quantified by qPCR. N=4 biological replicates. Mean±SD. One-way ANOVA. P-values are indicated.

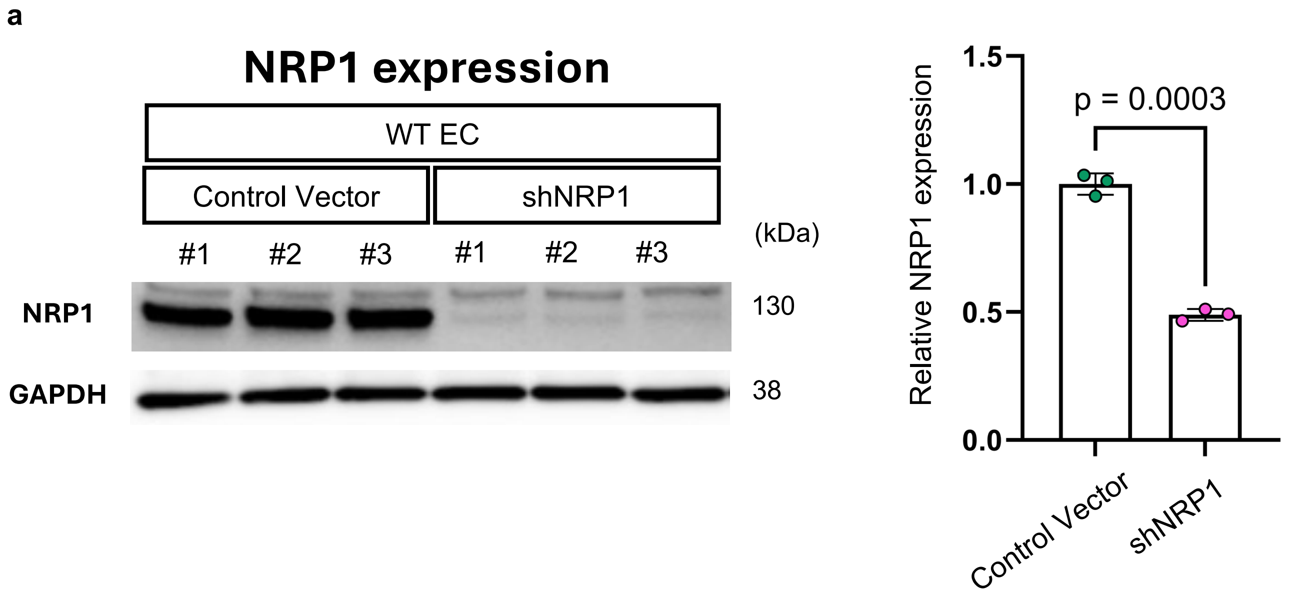


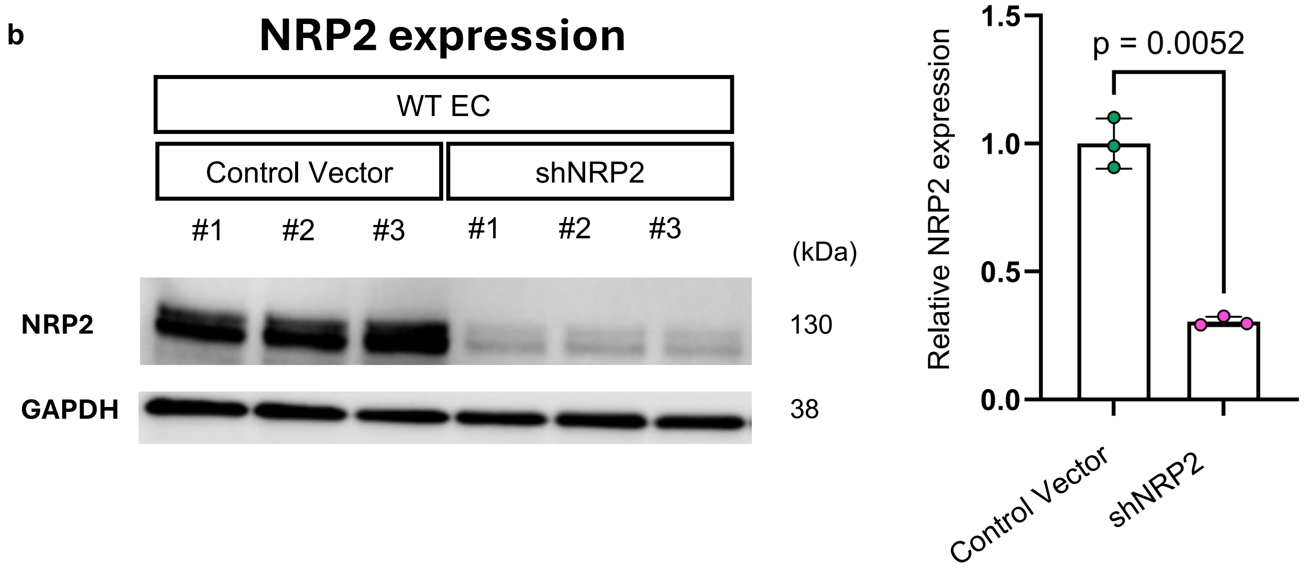


**Supplemental Figure 10. sh-RNA mediated knockdown of NRP1 and NRP2 in WT EC.** (a) Immunoblot of (a) NRP1 expression in WT EC that were treated with either control Vector or shNRP1 or (b) NRP2 expression in WT cells that were treated with either control Vector or shNRP2. Immunoblots were quantified (right panel). N=3 biological replicates. Mean±SD, Welch's t-test. P values are indicated.

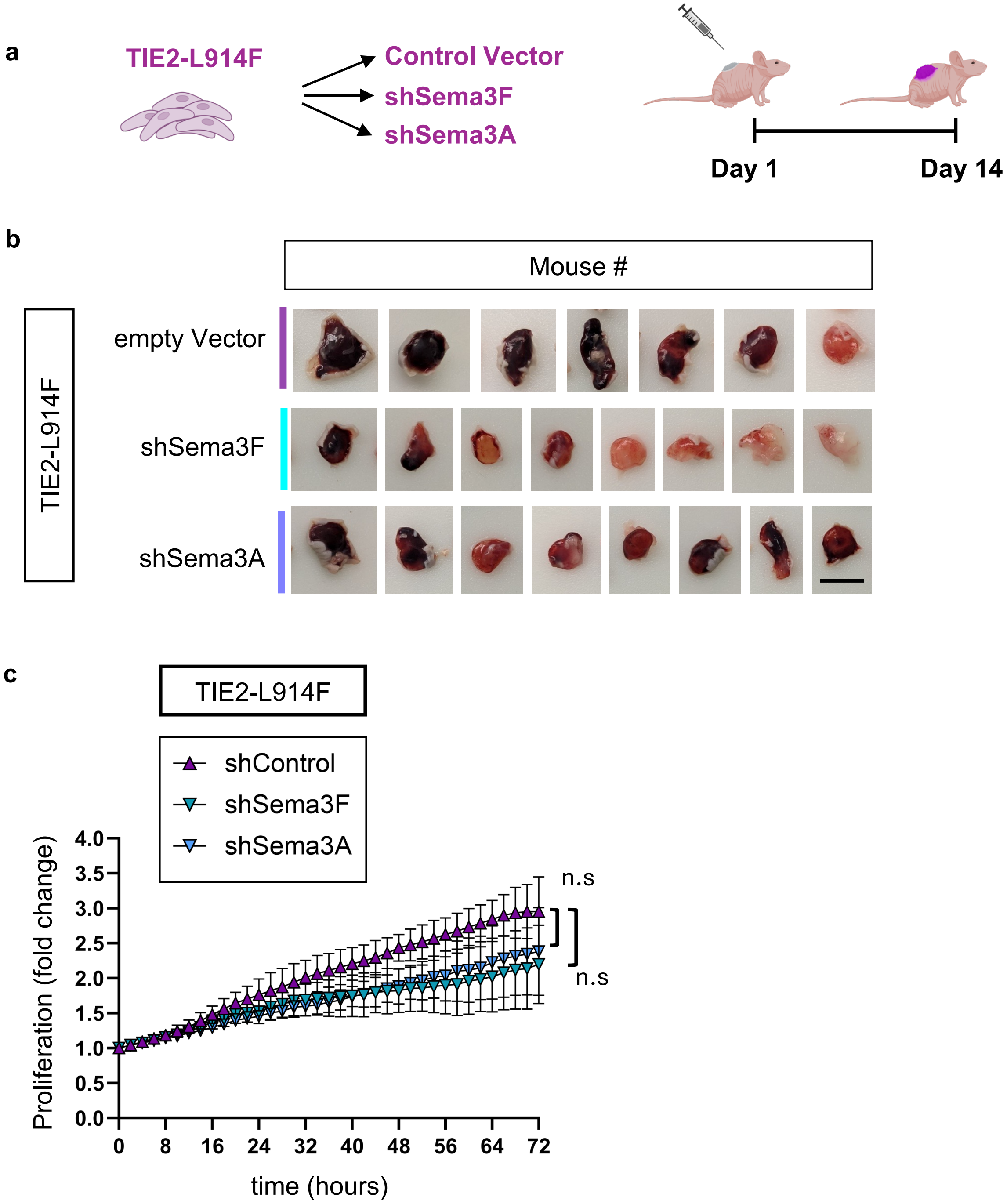


**Supplemental Figure 11**. **Sema3A and Sema3F exhibit cell autonomous effects on TIE2-mutant EC.** (a) Schematic of VM xenograft model with TIE2-L914F EC treated with control Vector or shRNA for Sema3F or Sema3A. Schematic created with Biorender.com (b) Photographs of the VM xenograft explants at day 14. n≥7 mice/condition. Scale bar: 1cm. (c) Proliferation analysis of TIE2 L914F EC treated with either control Vector, shSema3A or shSema3F. Mean±SD. n=12 wells/conditions in two independent experiments. 2-way ANOVA.

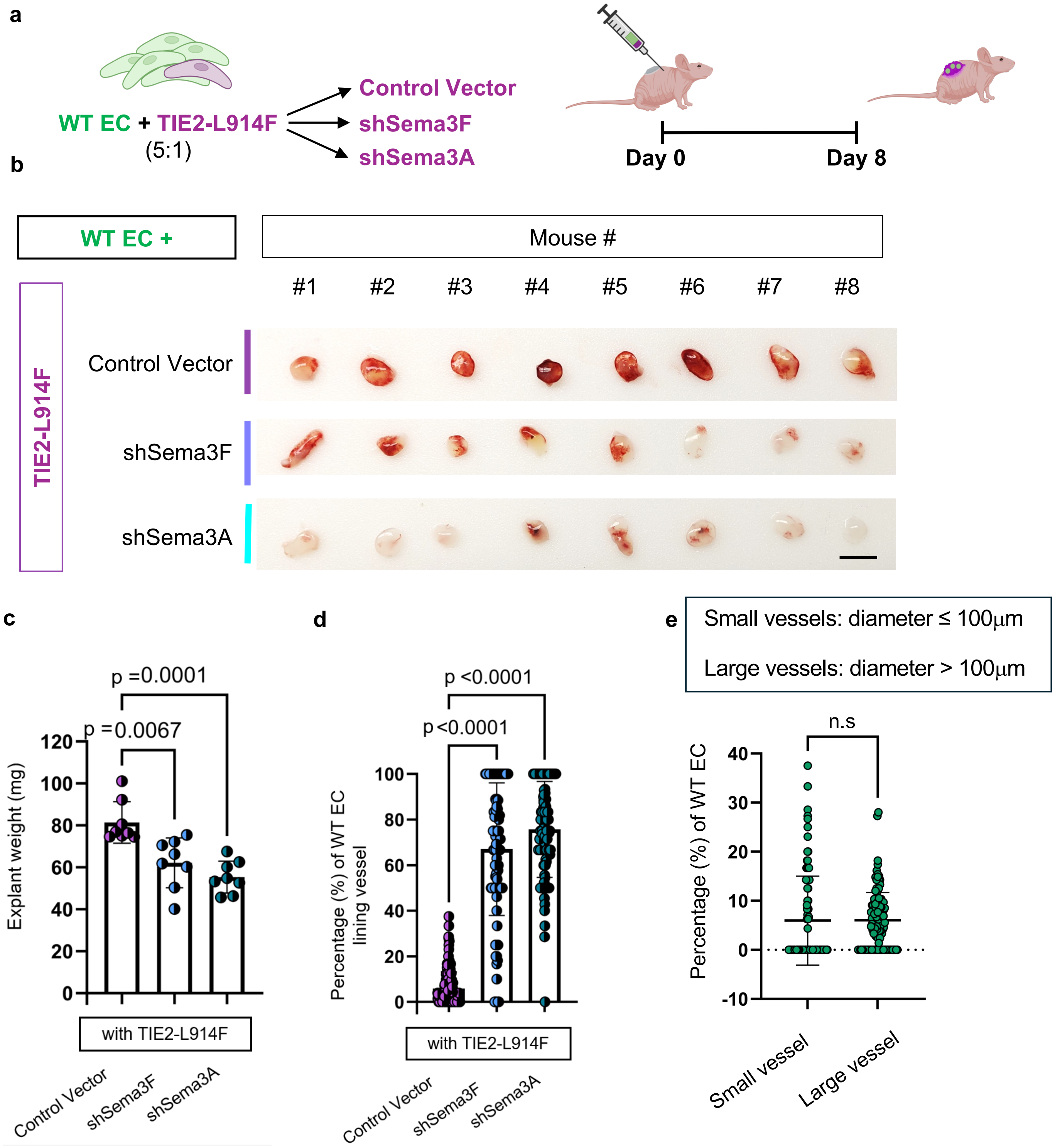


**Supplemental Figure 12. Knock-down of Sema 3A or 3F in TIE2-mutant EC restores cell communication with wild-type EC and reduces vascular lesion size.** (a) Schematic of the xenograft mixed EC model. Wild-type EC intermixed with HUVEC TIE2-L914F (empty Vector, shSema3A, or shSema3F), dissected on day 8. Schematic was created with Biorender.com. (b) Photographs of xenograft explants Scale bar: 1cm. (c) explant weights. (d) Quantification of cell type lining vessels. (e) Quantification of the percentage of wild-type EC in small vessels (≤100 μM) and large vessels (>100 μM). n=8 mice/condition and n≥115 vessels/condition, mean±SD, One-way ANOVA. p- values are indicated.
